# Differences in factors determining taxon-based and trait-based community structures: a field test using zooplankton

**DOI:** 10.1101/2024.04.12.589151

**Authors:** Hiromichi Suzuki, Hidetaka Ichiyanagi, Jamie M. Kass, Jotaro Urabe

**Author notes:** …. Corresponding author.

## Abstract

Ecological community structure, which has traditionally been described in terms of taxonomic units, is driven by a combination of dispersal and environmental filters. Traits have recently been recognized as alternative units for quantifying community parameters, but they may have important differences with taxonomic units. For example, as taxon-based community structures are determined by the identities of individual species, they may be more limited by species’ dispersal constraints, whereas trait-based community structures may be more regulated by environmental conditions because functional traits should have stronger linkages to habitat types rather than their specific locations. This suggests that the relative importance of dispersal and environmental filters may vary depending on the units employed to define community structure, yet how the contributions of these filters compare remains unclear. Zooplankton in lakes and reservoirs are an ideal system for examining the relative importance of these filters as they have moderate dispersal ability and inhabit communities in isolated habitats. In this study, we examined zooplankton assemblages in 87 artificial reservoirs throughout the Japanese archipelago to test the hypothesis that environmental filters play a more dominant role in determining trait-based community structures than taxon-based structures. To describe trait-based communities, we categorized zooplankton taxa into 12 functional groups using ecologically meaningful traits, including body size, feeding habits and mode, and population growth rate. We then examined the effects of both spatial configuration (reflecting dispersal filters) and environmental variables such as reservoir size and depth, trophic conditions, and fish assemblages for taxon- and trait-based community structures. Although variation in the taxon-based structure was explained equally well by spatial and environmental variables, variation in the trait-based structure explained by environmental variables was nearly twice that of spatial variables. These results support the idea that environmental filters play a more central role in determining trait-based community structures, and show that the relative importance of spatial and environmental filters changes with the way we define community structure.

**Open research statement:** All the data used in this study was obtained from disclosed sources including the Database of Dams in Japan (http://mudam.nilim.go.jp/home) for the location and watershed areas; the Japan Dam Foundation (http://damnet.or.jp/) for the dam specification data; and the River Environmental Database (Red: http://www.nilim.go.jp/lab/fbg/ksnkankyo/index.html) for water chemistry, fish, and zooplankton. Data obtained from local reports of the national reservoir management office, Japan, can be provided by the authors upon request. All analyses were conducted in the R programming language and all functions used in the present study originate from existing packages.

## Introduction

The structure of an ecological community, defined as a biological assemblage in a discrete habitat, is determined by a combination of filters, including those for species dispersal and the environment (Bilton et al., 2001; Palmer et al., 1996; Cottenie et al., 2001). Success of dispersal into a habitat patch is a function of the migratory abilities of the propagules, the distance from other habitats the species has already colonized, and the presence of geographic barriers between the habitats (Lockwood et al., 2005; Heino et al., 2015). For a species that has successfully passed the dispersal filter, its colonization success then depends on whether or not the local environmental conditions are a subset of its realized niche. These conditions can be defined by physical and chemical variables, such as habitat complexity, temperature, and nutrient supply (Schuler et al., 2017; Pearman et al., 2008), but also by biological variables, such as food resources, predators, and competitors (Dodson et al., 2000; Louette & De Meester, 2007; Hessen et al., 2006). Although these two filters are conceptually well understood, their relative importance in determining local community structure remains less so, and this is further complicated by the fact that community structure can be defined in different ways.

Community structure is commonly described using two different units: taxon-based ones that describe the community in terms of species composition, and trait-based ones that use ecologically meaningful functional traits, such as body size and feeding habits. Taxon-based units are essential for understanding the number and diversity of species in a community (Hutchinson, 1959; Tilman, 1982; Sax & Gaines, 2003), while trait-based units can help us to understand how communities are maintained under particular environments (Lindeman, 1942; Simberloff & Dayan, 1991; Cadotte et al., 2011; Louca et al., 2016; Pomerleau et al., 2015). Furthermore, the factors most important for regulating community structure may differ depending on how the structure itself is quantified. On one hand, similarity in community composition should decrease with increasing habitat distance (Nekola & White, 1999) as species have different dispersal capacities over limited spatial scales that lead to different species pools among habitat patches. Thus, dispersal filters should play a substantial role in determining taxon-based community structure. On the other hand, the niche space of a habitat is structured by local physical, chemical, and biological conditions that may be similar among habitats regardless of distance. In the case that different habitats share available niche space but differ in species composition, these species are likely to be similar in functional traits (Messier et al., 2010). This suggests that, regardless of distance from other habitats, environmental filters should play a critical role in determining trait-based community structure. Therefore, it is hypothesized that environmental filters play a more important role in regulating trait-based community structures, while dispersal filters have a greater effect on taxon-based community structures. In partial support of this hypothesis, several studies have demonstrated the importance of environmental filters in determining the functional structure of animal communities (e.g., Loewen et al., 2019; Vogt et al. 2013). However, no studies have yet compared the relative importance of both filters in regulating different community structures, perhaps because this would necessitate the difficult task of comparing a large number of isolated communities at different distances.

A good model system for testing this hypothesis is freshwater zooplankton, which live in discrete aquatic habitats surrounded by land (i.e., lakes, ponds, and reservoirs) with distinct communities that can be easily defined (Hortal et al., 2014). These lentic habitats vary environmentally depending on their size, depth, and watershed characteristics (Arbuckle & Downing, 2001). Further, as lentic habitats are scattered across landscapes and zooplankton are easily collected via water samples, comparing communities established in different environmental conditions and locations is relatively straightforward. In addition, zooplankton represent various taxa, including rotifers, cladocerans, and copepods, and differ in body size, feeding habits, morphology, and behavior. Thus, zooplankton community structure can be described using both taxon-based and trait-based units.

In general, zooplankton have dormant stages that can tolerate harsh conditions, such as drought and low temperatures. These tolerances allow them to migrate and disperse propagules to distant habitats, for example, by drifting on floods (Bozelli et al., 2015), riding the wind (Pinceel et al., 2016), or hitchhiking on waterfowl (Green & Figuerola, 2005; Hessen et al., 2019), although this ability appears to vary widely among taxa (Jenkins & Underwood, 1998; Gray & Arnot, 2011). As such, zooplankton are ideal organisms to examine the relative importance of environmental and dispersal filters in determining taxon- and trait-based community structure.

In the present study, we analyzed the zooplankton communities in 87 reservoirs that span the wide latitudinal range of the Japanese archipelago. First, we examined the taxon-based and trait-based structures of the zooplankton communities in these reservoirs, then analyzed the dominant factors regulating these structures. In this analysis, we considered geographic locations of reservoirs at different scales as spatial factors, and dam specifications, water chemistry, and fish assemblages as environmental factors. Using the results of these analyses, we tested the hypothesis that the environmental filter plays a dominant role in regulating the trait-based community structure, while the dispersal filter has a greater effect on the taxon-based community structure (Fig. 1).

**Figure 1.**
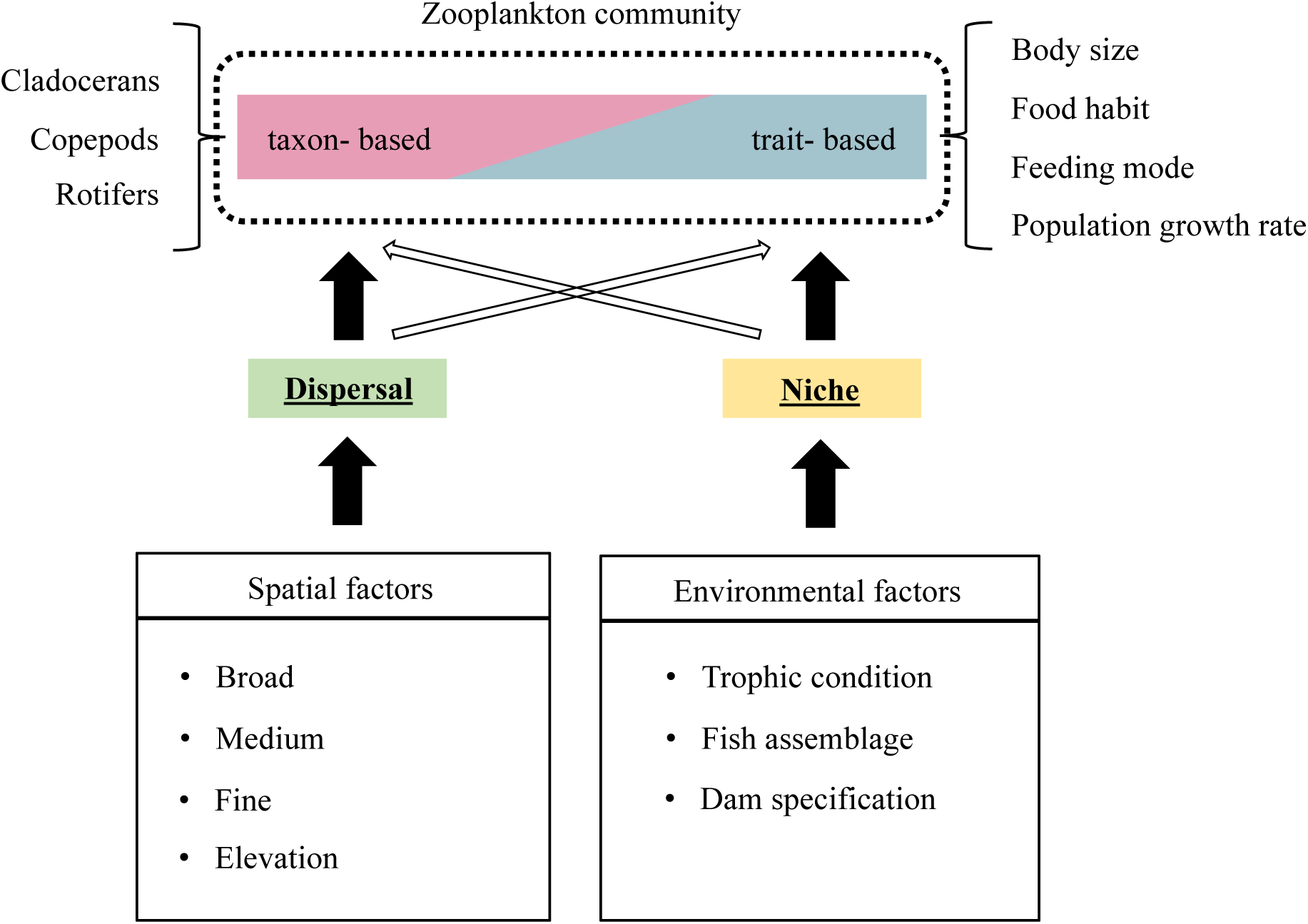
Conceptual diagram showing the workflow of our analysis. We quantified the relative importance of dispersal and niche filters by examining the relationship between taxon- or trait-based community structures and environmental or spatial factors.

## Materials and Methods

We examined zooplankton communities in 87 reservoirs located in a wide geographical area of the Japanese archipelago, from 26.48194° to 44.11527° north latitude, from 127.94944° to 143.38778° east longitude, and from 28 to 1123 m in elevation (Fig. 2). Zooplankton data have been routinely collected in these reservoirs by the National Census on River and Dam Environment, which has been conducted by the Ministry of Land, Infrastructure, Transport and Tourism (MLIT) of Japan every five years since 1990. In the program, surveys were carried out for one of the five years, with samples being taken 4-12 times per survey year. In this study, we analyzed the zooplankton data collected in each reservoir at seasonal (four times) and monthly (twelve times) scales for one year each during the 2001-2005 period (3^rd^ round of sampling) and the 2006-2010 period (4^th^ round of sampling). In 52 of the 87 reservoirs, zooplankton were collected in both the 3rd and 4th sampling rounds, but in the remaining reservoirs, zooplankton were collected in only one of these single rounds. Since the 3^rd^ and 4^th^ sampling rounds were five years apart and we used year-specific environmental data, we treated each sampling round independently, even for the same reservoirs. Thus, we obtained a total of 139 datasets of the zooplankton community.

**Figure 2.**
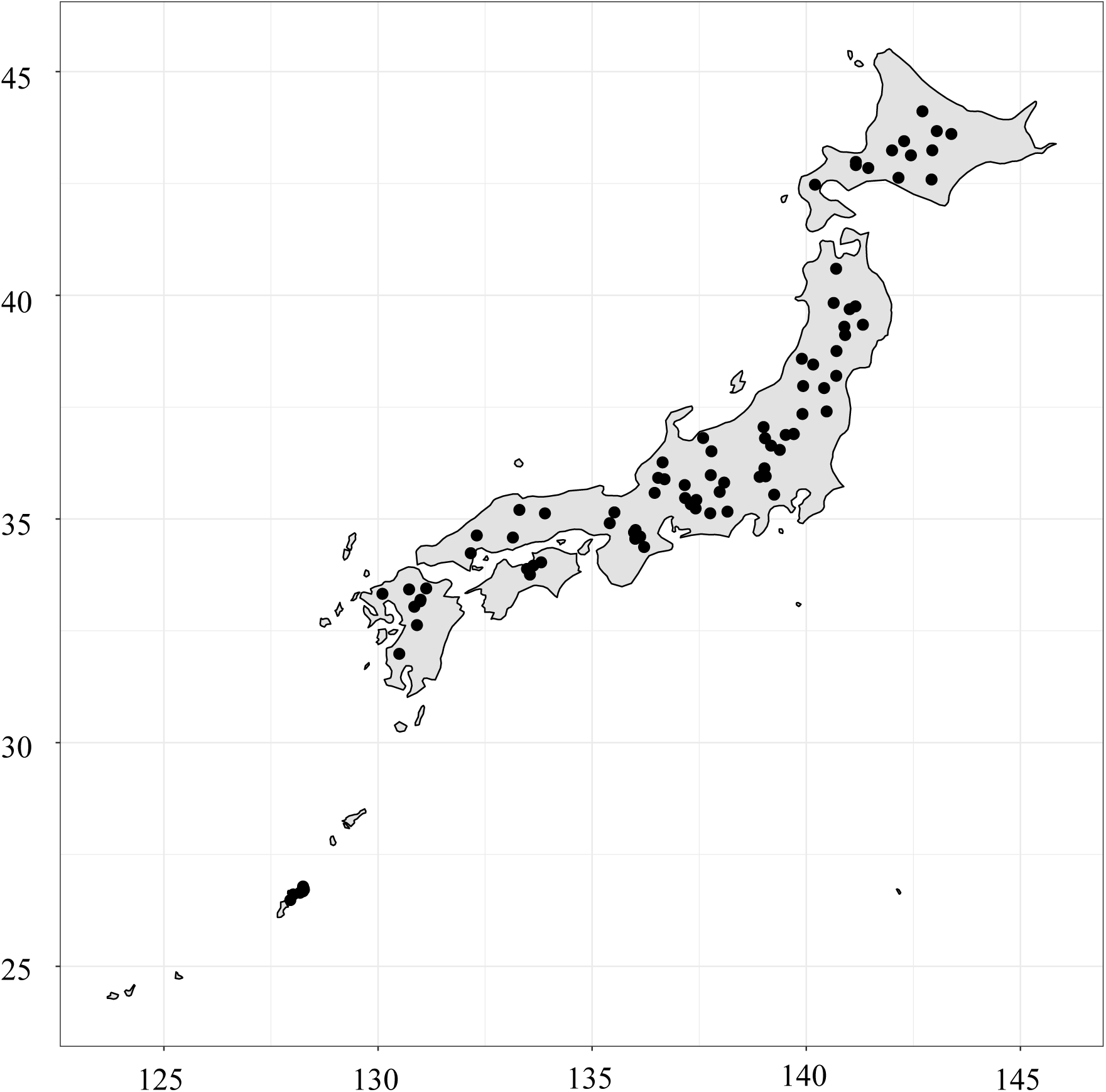
Map showing locations of artificial reservoirs in the Japanese archipelago from which we sampled zooplankton in this study.

In each reservoir, the zooplankton samples were collected at a site near the dam or at the deepest point by vertical tows of conical plankton nets with 100-µm mesh from the bottom or mid-depth to the surface. Collected zooplankton were preserved in Lugol’s solution or 4% formalin solution and examined under a microscope for identification. Specimens that could not be identified to the species level were classified to the genus level, except for calanoid copepods, which were often classified to the family or order level due to uncertainty in identification even for genera. As the volume of water passing through the plankton nets was different among reservoirs, we did not consider abundance data for zooplankton taxa, and instead reduced counts to binary presence/absence data.

### Taxa and trait units

For each reservoir, we assigned a score of 1 (presence) to the genus or species that occurred in at least one sample for that year, and a 0 (absence) when no samples were available within a year. We then estimated the frequency of occurrence for each species or genus across all the reservoirs and years and created a taxonomic list consisting of species or genera that occurred in more than 15% of the 139 zooplankton datasets. We used these binary datasets as taxon-based community structures.

To describe the zooplankton community using trait-based units, we first classified all zooplankton taxa based on body size as follows: large: > 1 mm, medium: 0.5-1 mm, and small: < 0.5 mm in total length excluding the apical spine and external appendage. Although these body-size categories are arbitrary, body size is known to be closely related to predator vulnerability (Zaret, 1980; Lampert & Sommer, 2007) and physiological rates, including metabolic (Ikeda 1985; Lampert 1984) and feeding rates (Peters & Downing 1984). Accordingly, it is often considered a “master trait” (Litchman & Klausmeier 2008; Litchman et al. 2013). We then divided each of the body-size groups further into several distinct groups according to trophic level (carnivore, omnivore, herbivore, detriti-herbivore, and detritivore), feeding mode (filter feeders, raptorial feeders, and browsers) and population growth rate. Here, “detriti-herbivore” refers to the diet of animals that feed on both suspended algae and sedimented detritus. Trophic level was classified based on previous knowledge (Benzie, 2005; Barnett et al., 2007; Branstator, 2005; Rizo et al., 2017; Kuczyńska-Kippen et al., 2020; Dussart & Defaye, 2001; Koste & Shiel, 1991; Duggan, 2001; Gilbert, 2022). Feeding mode, or the method of resource acquisition, was classified according to Sommer & Stibor (2002), Branstator (2005) and Barnett et al. (2007) for cladocerans; Dussart & Defaye (2001) for copepods; and Hillbricht-Ilkowska (1983) and Gilbert (2022) for rotifers. For some rotifer taxa, no published information regarding feeding mode was available; in that case, feeding mode was determined based on the morphology of the trophis (i.e., the grinding apparatus) according to Oh et al. (2017) and Palazzo et al. (2021). Groups with different maximum population growth rates were classified according to Allan (1976). Thus, using body size, feeding habits, feeding mode and population growth rate, we classified the zooplankton taxa into 12 functional groups (Table 1, see Table S2 for all taxa). For each of the 139 zooplankton community records, we assigned a score of 1 (presence) to the functional group in which at least one taxon occurred, and a 0 (absence) when this group was not represented. The functional groups present in each community were then used to describe the trait-based community structure.

**Table 1.** Functional groups used for description of the trait-based community.

| Functional category | Body size | Feeding mode | Food habit | Population growth rate |
| --- | --- | --- | --- | --- |
| Large raptorial-feeding carnivore | Large | Raptorial | Carnivore | Middle |
| Large filter-feeding herbivore | Large | Filter | Herbivore | Middle |
| Large browsing omnivore | Large | Browser | Omnivore | Low |
| Large filter-feeding omnivore | Large | Filter | Omnivore | Low |
| Medium filter-feeding herbivore | Medium | Filter | Herbivore | Middle |
| Medium raptorial-feeding omnivore | Medium | Raptorial | Omnivore | Low |
| Medium raptorial-feeding carnivore | Medium | Raptorial | Carnivore | High |
| Small filter-feeding herbivore | Small | Filter | Herbivore | Middle |
| Small filter-feeding detriti-herbivore | Small | Filter | Detritivore /<br>Herbivore | Middle |
| Small raptorial-feeding herbivore | Small | Raptorial | Herbivore | High |
| Small raptorial-feeding omnivore | Small | Raptorial | Omnivore | High |
| Small filter-feeding detritivore | Small | Filter | Detritivore | High |

### Spatial and environmental variables

The elevation, latitude, and longitude of the reservoirs were obtained from the Database of Dams in Japan (http://mudam.nilim.go.jp/home). In addition, data specifying dam characteristics, including watershed area (Watershed), effective storage volume (Volume), dam height (Height), dam width (Width), and years elapsed since the reservoir first impounded its full capacity after dam construction (Age) were obtained from the database published by the Japan Dam Foundation (http://damnet.or.jp/Dambinran/binran/) and the periodic reports issued by each reservoir management office. In some reservoirs, a vertical water aeration-based circulation facility (VWCF) was installed to reduce algal production by moving them to deeper depths. As algae are important food resources for many zooplankton, such facilities may affect community structures. We therefore included the presence or absence of a VWCF as an environmental factor. Water retention time of the reservoir was estimated by dividing the effective water storage volume by the annual inflow rate in that year, which was obtained from the Database of Dams in Japan. For reservoirs with no data available from the database, we obtained the annual inflow data directly from the periodic reports issued by the reservoir management office.

Water chemistry data relating to total phosphorus (TP), total nitrogen (TN), and chlorophyll *a* (Chl-*a*) measured monthly in the reservoir were obtained from the River Environmental Database (RED http://www.nilim.go.jp/lab/fbg/ksnkankyo/index.html) established by MLIT. For the analyses described below, we used the mean annual values of these variables for the year in which the zooplankton were collected. From the RED, for each site we also obtained presence/absence records of fish species (total 84) distributed in each reservoir during the period of the 3^rd^ or 4^th^ round of collection (depending on the record).

### Statistical analyses

As above, we analyzed a total of 139 zooplankton communities obtained from 87 reservoirs. All statistical analyses were performed using R version 4.0.4 (R core team 2021). Spatial configurations among the reservoirs were modeled using distance-based Moran’s eigenvector maps (dbMEM) based on the geographic coordinates of the reservoirs using the function “*dbmem”* in the package *adespatial* (Dray et al., 2022). Moran’s eigenvalues were estimated from the geographic distance matrix (latitude and longitude) and used as variables representing the degree of spatial autocorrelation at different scales between study sites, but also the degree of dispersal limitation of zooplankton communities. The significance of each Moran’s eigenvalue was tested using the function “*moran.randtest”* in the R package *adespatial*. We labeled the eigenvalues in order from the largest to the smallest spatial scale, using the notation MEM 1, MEM 2, and so on.

We summarized the fish assemblages in the reservoirs using a principal coordination analysis (PCoA) based on the Sorensen dissimilarity matrix using the function *“cmdscale”* in the R package *stats* (R Core Team, 2021). Before running the analysis, we removed fish species that occurred in less than 15% of the 139 species lists to minimize the effect of rare species on the characterization of the fish assemblage. This criterion was arbitrary, but sensitivity tests for 5% and 10% showed that results were not largely changed. For zooplankton analyses, we used the scores of the first three axes of the PCoA as fish assemblage data (list the percentages of variance explained for each axis). In addition, to characterize each axis we calculated species scores for each axis using the function “*add.spec.score*” in the R package *BiodiversityR* (Kindt & Coe, 2005).

To determine the ability of the environmental and spatial filters to explain each of the taxon- and trait-based community structures, we performed a partial distance-based redundancy analysis (partial-dbRDA). This analysis is an extension of multiple regression for community data that has been often used to compare between different candidate explanatory variable-sets (Legendre & Legendre, 2012). For environmental filter variables, we used the variables listed in Table 2 and fish assemblage PCs 1–3. For spatial filter variables, we used MEM 1–12 and elevation. All variables besides the PCoA scores and Moran’s eigenvalues were prior to analysis. We calculated variance inflation factors among these explanatory variables using function “*cor*” in the R package *stats,* and “*solve*” and “*diag*” in the R package base (R core team, 2021). These were less than 10, indicating negligible variable collinearity (O’Brien 2007). To perform patial-dbRDA, we first estimated the Sorensen dissimilarity index of the zooplankton composition between the reservoirs using the function “*capscale*” in the R package *vegan* (Oksanen et al., 2020). To distinguish the effects of environmental and spatial filter variables, one model was constructed in the partial-dbRDA to focus on each variable and adjust for the other. For these models, we tested the statistical significance of the focal variables using permutation tests (n = 1999) with the function “*anova.cca”* in the R package *vegan*. If the focal variables were significant, we performed variation partitioning using the function “*varpart*” in R the package *vegan*. This procedure helped us to examine the relative importance of the different filter variables in determining the taxon- and trait-based community structures.

**Table 2.** Summary of environmental variables showing maximum, minimum, mean, median, and standard deviation (SD).

| Variables | Min | Max | Mean | Median | SD |
| --- | --- | --- | --- | --- | --- |
| **Nutrient condition** |  |  |  |  |  |
| TP (mg/l) | 0.003 | 0.09 | 0.017 | 0.012 | 0.149 |
| TN (mg/l) | 0.100 | 1.511 | 0.449 | 0.361 | 0.279 |
| Chl *a* (mg/m<sup>3</sup>) | 0.004 | 35.000 | 6.351 | 4.113 | 6.266 |
| **Dam specifications** |  |  |  |  |  |
| Watershed (km<sup>2</sup>) | 7.4 | 2409 | 342.5 | 225.5 | 409.5 |
| Volume (1000m<sup>3</sup>) | 1250 | 289000 | 55960.1 | 42000 | 52427 |
| Height (m) | 24 | 156 | 89.06 | 86 | 31.3 |
| Width (m) | 127.6 | 1480 | 340.2 | 322 | 173 |
| VWCF | 0 | 1 | - | - | - |
| Turnover (/year) | 1.106 | 922.140 | 29.025 | 7.546 | 92.127 |
| Age (years) | 2 | 54 | 25.91 | 26 | 13 |

As the above analysis included multiple data records that originated from repeat surveys within the same reservoirs, which might bias our results in favor of conditions at sites with multiple survey records, we performed a randomization test to determine if and to what extent our results changed if repeat surveys were excluded. For the reservoirs with records from both the 3rd and 4th round surveys (n=52), we first randomly selected one of these two survey records from each reservoir. Together with the data from reservoirs with only one survey (3rd or 4th round; n=25), we created a rarefied dataset of individual records from each of the 87 reservoirs. We then performed the same analysis as above using this rarefied dataset and estimated the difference in the percentage of variance explained by spatial and environmental variables. We repeated this procedure 1000 times. With this analysis, we tested the robustness of our results using all data from all reservoirs.

Finally, we examined the effects of each individual explanatory variable on community structure using partial-dbRDA to determine the relative importance of these variables in determining the community structure. In this analysis, the target variable in each partial-dbRDA model was treated as the objective variable and the other variables as “conditions”. We tested the statistical significance of each explanatory variable using the permutation tests as described above.

## Results

### Environmental conditions of reservoirs

The reservoirs varied widely in effective storage volume (Arakawa: 1,250,000 m^3^ to Sameura: 289,000,000 m^3^), dam height (Ikeda: 24 m to Miyagase: 156 m), watershed area (Arakawa: 7.4 km^2^ to Maruyama: 2409 km^2^), and age (Ishibuchi: 54 years to Origawa: 2 years [from full capacity when zooplankton were first sampled]) (Table 2). Of the 87 reservoirs sampled, 13 had at least one VWCF. Nutrient status varied among the reservoirs from oligotrophic to eutrophic as TP, TN, and chlorophyll-*a* concentrations ranged from 0.003 to 0.09 mg L^-1^ (median 0.012 mg L^-1^), 0.1 to 1.511 mg/L (0.361 mg L^-1^), and 0.004 to 35 µg L^-1^ (4.113 mg L^-1^), respectively.

### Spatial configurations of reservoirs

We obtained 12 statistically significant Moran’s eigenvalues for the reservoirs, which we refer to as MEM 1 through MEM 12 (Table S1; Fig S1). Some of these eigenvalues are visualized as examples in Figure 3, and maps of all 12 significant Moran’s eigenvalues are shown in Figure S1. The number of spatial groups ranged from 2 (the largest scale) to 20 (the smallest scale) (Fig. 3).

**Figure 3.**
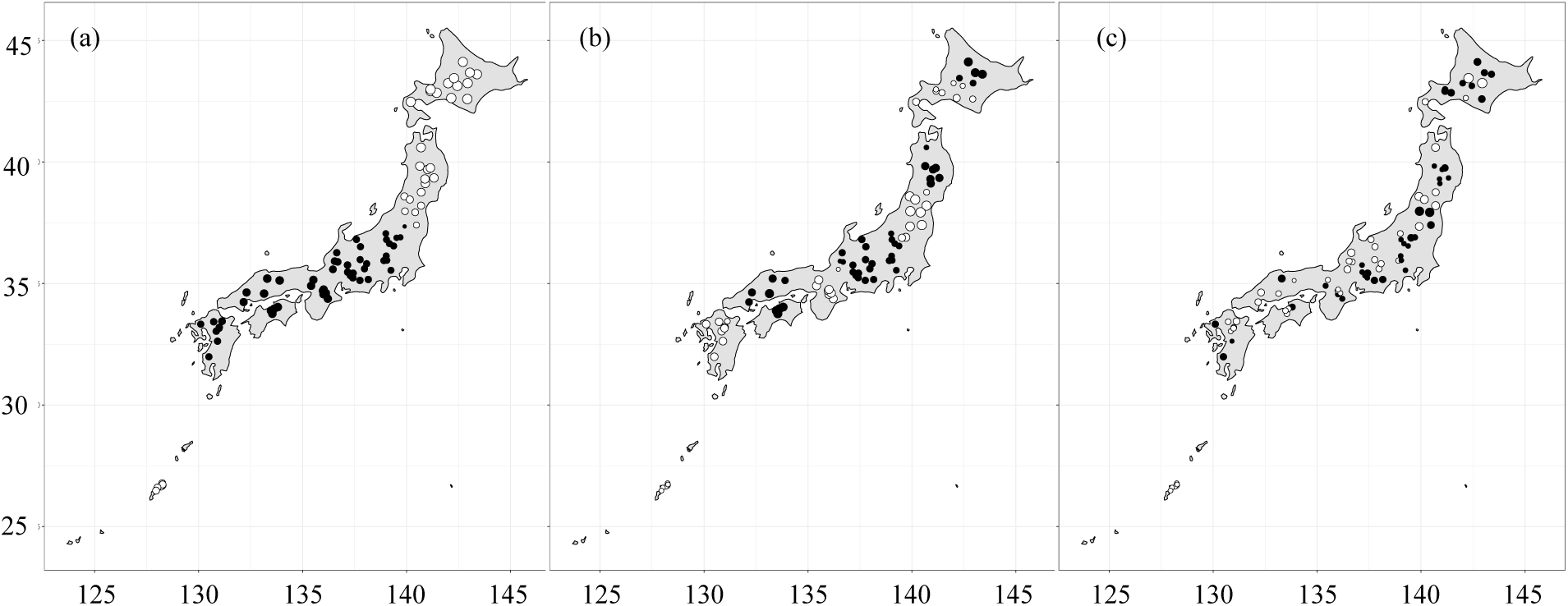
Maps showing examples of Moran’s eigenvalues that we calculated for reservoir locations and used as spatial variables for dispersal filters in our analysis. The largest spatial scale is represented by MEM 1 (a) and scales become smaller with increasing numbers: MEM 4 represents a medium scale (b), while MEM 11 represents a small scale (c). The white and black circles represent positive and negative eigenvalues, respectively, and the size of the circle represents the magnitude of the absolute value.

### Zooplankton

In a total of 139 lists of zooplankton community composition obtained from one- or two-year surveys for the 87 reservoirs, we identified 131 taxa, including 33 cladocerans, 29 copepods, and 69 rotifers (Table S2). The reservoir with the highest number of taxa per year was Origawa with 52, while the one with the lowest was Omachi with 5, and the overall average among reservoirs per year was 21 taxa. Of these, 38 taxa were recorded in more than 15% of the zooplankton lists (Table S2).

We then assigned the 131 taxa to the 12 functional groups based on body size, feeding habits, feeding mode, and maximum growth rate (Table S2). The reservoir with the highest number of functional groups per year was Hitokura with 11, while that with the lowest was Maruyama with 4, and the average 8.2 groups per reservoir per year. The Sorensen index ranged from 0 to 1 with a mean of 0.559 for the taxon-based community and from 0 to 0.828 with a mean of 0.202 for the trait-based community (Fig. S2). Because the number of units describing community structure was smaller for the trait-based community (12 functional groups) than for the taxon-based community (38 taxa), the coefficient of variation (CV) of the Sorensen dissimilarity index was smaller for the taxon-based communities (30.3%) than for the trait-based community (59.5%).

### Fish assemblages

The PCoA for fish communities explained a total of 32.0% of the variation in community structure among reservoirs with the first three axes (Tables 3 and S3). For PC1, warm-water fishes, such as *Pseudogobio ecocinus ecosinus* (pike gudgeon), *Micropterus salmoides salmoides* (largemouth bass)*, Zacco platypus* (pale chub), and *Rhinogobius* sp. (goby) were positively related, whereas cold-water fishes, such as *Salvelinus* sp. (char) and *Oncorhynchus mykiss* (rainbow trout) were negatively related. *M. salmoides salmoides* is piscivorous whereas *P. ecocinus ecosinus* and *Rhinogobius* sp. are omnivorous and prey on benthic animals. Therefore, we refer to PC1 as the warm-water euryphagous fish assemblage (WEF). For PC2, *Leuciscus hakonensis / ezoensis* (Japanese dace), *Hemibarbus labeo barbus (barbel steed)*, *Opsariichthys uncirostris uncirostris* (three-lips), *Squalidus* sp. (gudgeon), and *Zazzo platypus* showed relatively large positive eigenvectors (Table 3; Table S3). Since all these fishes were cyprinids, we refer to PC2 as the cyprinid fish assemblage (CYF). For PC3, *Hypomesus nipponensis* (pond smelt) was positively associated with the highest loading (Table S3). This fish species is planktivorous and has been artificially stocked in various lakes and reservoirs for commercial and sport fishing. Therefore, we refer to PC3 as the smelt-dominated fish assemblage (SDF).

**Table 3.** Results of the principal coordination analysis showing the eigenvalue, viability, and cumulative variability of the first three coordinates, with the top 5 fish taxa having the highest absolute eigenvector value in these coordination axes. The direction of the eigenvector is shown in parentheses.

| Axis | PC1 | PC2 | PC3 |
| --- | --- | --- | --- |
| **Eigenvalue** | 33.04 | 15.86 | 11.32 |
| **Cumulative variability (%)** | 17.58 | 26.02 | 32.04 |
| Rank |  |  |  |
| 1 | *Zacco platypus* (+) | *Zacco platypus* (+) | *Hypomesus nipponiensis* (+) |
| 2 | *Salvelinus* sp. (-) | *Leuciscus hakonensis* /<br>*ezoensis* (+) | *Oncorhynchus masou masou*<br>(+) |
| 3 | *Pseudogobio ecocinus ecocinus*<br>(+) | *Hemibarbus labeo barbus* (+) | *Phoxinus* sp. (+) |
| 4 | *Rhinogobius* sp. (+) | *Squalidus* sp. (+) | *Misgurnus anguillicaudatus*<br>(+) |
| 5 | *Micropterus salmoides*<br>*salmoides* (+) | *Opsariichthys uncirostris*<br>*uncirostris* (+) | *Cyprinus carpio* (+) |
| Name of fish assemblage | Warm-water euryphagous fish<br>assemblage (WEF) | Cyprinid fish assemblage<br>(CYF) | Smelt-dominated fish<br>assemblage<br>(SDF) |

### Relative importance of environmental and spatial filters

Partial-dbRDA showed that both environmental and spatial filter variables significantly explained the variation in both the taxon-based and trait-based structures of zooplankton communities (Table S4). The variation partitioning analysis showed that about one-third of the total variation in each community structure was explained by these variables, with different relative proportions for the different variable types. Of the 29.0% total variance explained for the taxon-based structure, the unique proportions for environmental (10.9%, 18.8% with covariance) and spatial (10.2%, 18.1% with covariance) variables were similar (Fig. 4; Table S5). On the other hand, of the 39.2% total variance that was explained for the trait-based structure, the unique proportion for environmental variables (19.9%, 27% with covariance) was 1.63 times higher than that for the spatial variables (12.2%, 19.3% with covariance) (Fig. 4; Table S5). We obtained the same results for the rarefied datasets created for the randomization test: the difference in the total variance explained between environmental and spatial variables was always larger for trait-based community than taxon-based community in 99.9% of the 1000 rarefied datasets (Fig. S3).

**Figure 4.**
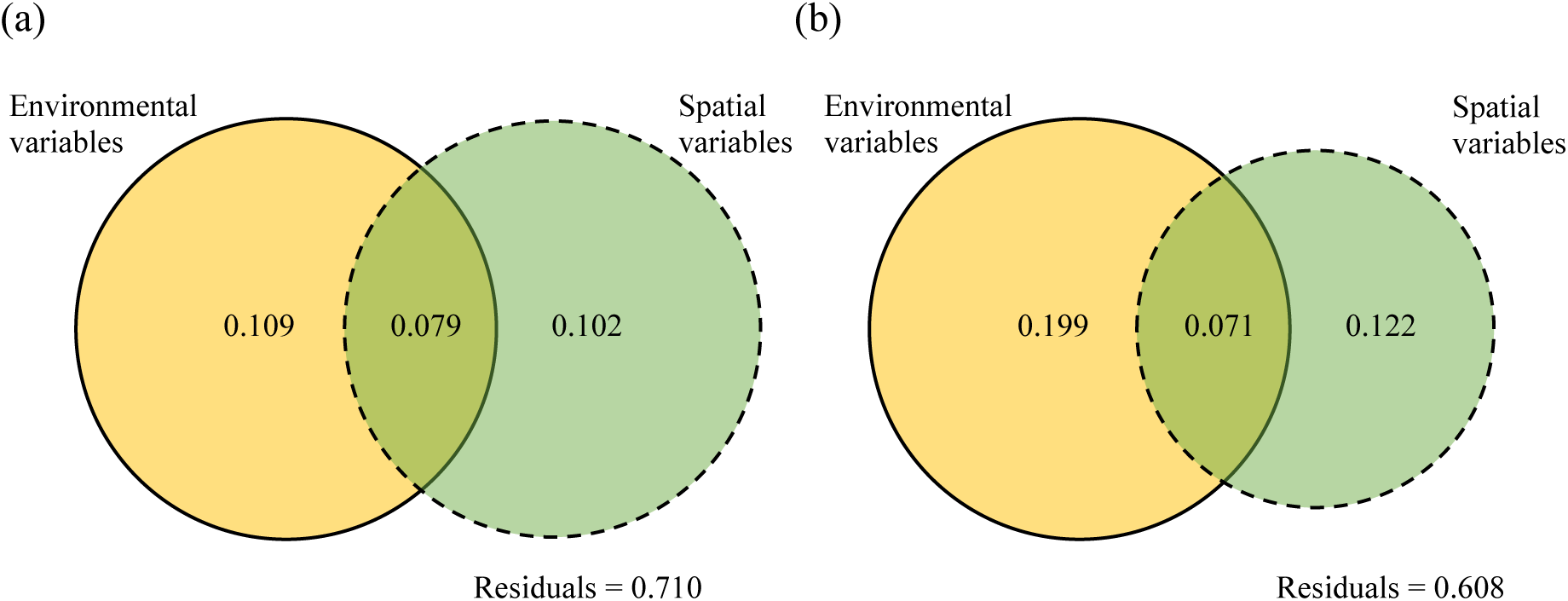
Venn diagrams showing the contributions of environmental variables, spatial variables, and residuals to the total variation in the taxon-based (a) and trait-based (b) zooplankton community structures.

### Effects of individual variables on the taxon-based community

The partial-dbRDA showed that while dam characteristics (watershed area, reservoir water volume and dam height) and fish communities [WEF (PC 1) and SDF (PC 3)] had significant effects on taxon-based community structure (Table 4), lake trophic conditions (TP, TN, and Chl-*a*) did not. The highest percentages of contribution to the variation (adjusted R^2^) were watershed area and dam height, followed by water volume and fish communities WEF and SDF. Specifically, rotifer species such as *Ascomorpha* spp., *Asplanchna priodonta*, and *Ploesoma truncatum* were positively associated with reservoir water volume and SDF, while the calanoid copepods and *Daphnia galeata* were positively associated with dam height and WEF (Table S6). In contrast, *Cyclops vicinus*, *Alona* spp. and *Asplanchna priodonta* were negatively associated with WEF (Table S6). The presence of VWCF had no significant effect on the variation in taxon-based community structure among reservoirs.

**Table 4.** Results of partial-dbRDA for the taxon-based community showing adjusted coefficients of determination for environmental and spatial variables with significant probability. For significant variables, the direction of the effects is shown in parentheses. Bold type indicates the variables that were significant.

| Environment variables | adjusted R<sup>2</sup> | F | *p* | Spatial variables | adjusted R<sup>2</sup> | F | *p* |
| --- | --- | --- | --- | --- | --- | --- | --- |
| TP | 0.011 | 1.510 | 0.067 | Elevation (+) | **0.020** | **2.300** | **0.004** |
| TN | 0.006 | 1.041 | 0.395 | MEM1 | 0.007 | 1.100 | 0.324 |
| Chl *a* | 0.004 | 0.843 | 0.65 | **MEM2 (+)** | **0.020** | **2.308** | **0.003** |
| **WEF (PC1) (+)** | **0.017** | **2.085** | **0.004** | **MEM3 (+)** | **0.0201** | **2.402** | **0.002** |
| CYF (PC2) | 0.005 | 0.955 | 0.515 | **MEM4 (+)** | **0.015** | **1.852** | **0.012** |
| **SDF (PC3) (-)** | **0.013** | **1.661** | **0.034** | **MEM5 (+)** | **0.016** | **1.941** | **0.014** |
| **Watershed (+)** | **0.036** | **3.755** | **< 0.001** | MEM6 | 0.010 | 1.378 | 0.123 |
| **Volume (+)** | **0.025** | **2.766** | **< 0.001** | MEM7 | 0.010 | 1.407 | 0.106 |
| **Height (-)** | **0.029** | **3.166** | **< 0.001** | MEM8 | 0.010 | 1.405 | 0.109 |
| Width | 0.003 | 0.757 | 0.766 | **MEM9 (+)** | **0.022** | **2.483** | **< 0.001** |
| VWCF | 0.005 | 0.986 | 0.468 | **MEM10 (-)** | **0.014** | **1.742** | **0.025** |
| Turnover rate | 0.004 | 0.809 | 0.638 | **MEM11 (+)** | **0.015** | **1.825** | **0.02** |
| Age | 0.007 | 1.135 | 0.289 | MEM12 | 0.003 | 0.732 | 0.777 |

Among the spatial variables, elevation and large- and small-scale MEMs significantly affected the taxon-based community structure (Table 4). Although the percentage contributions of these variables to the total variation were relatively small (1.4% to 2.2%), the number of significant variables affecting taxon-based community structure was greater for spatial factors than for environmental ones. Among the spatial variables, the cladocera *Daphnia dentifera* was positively associated with elevation and MEM 10, while the copepod *C. vicinus* was associated negatively with elevation and MEM 4 (Table S7). Similarly, different rotifer species were associated with different MEM variables. Some cladocerans, such as *Bosminopsis deitersi*, *Daphnia galeata* and *D. dentifera*, showed high eigenvector values for small-scale MEMs (Table S7). In contrast, some rotifers such as *Ascomorpha ovalis, Cephalodella* sp. and *Collotheca* spp. showed high eigenvector values for larger-scale MEMs (MEM 2 to 5).

### Effects of individual variables on the trait-based community

For the trait-based community structure, the environmental variables for fish assemblages WEF and CYF, as well as reservoir age, watershed area, water volume, and dam height had significant effects (Table 5). The highest percentages of contribution to variation (adjusted R^2^) were WEF, CYF, and reservoir age (Table 5). Similar to the taxon-based community, TP, TN, Chl-*a*, and the presence of VWCF had no significant effect on this community structure. Among functional groups, large filter-feeding herbivores and omnivores were positively associated with WEF and negatively associated with watershed area (Table S8). In addition, water volume had positive associations with large raptorial-feeding carnivores and medium and small filter-feeding herbivores, yet negative ones with small filter-feeding detriti-herbivores and small raptorial-feeding herbivores. Large filter feeding omnivores and small filter-feeding herbivores were associated with reservoir age (Table S8). Large filter-feeding omnivores and small raptorial-feeding herbivores were positively associated with CYF (Table S8).

**Table 5.** Results of partial-dbRDA for the trait-based community showing adjusted coefficients of determination for environmental and spatial variables with significant probability. For significant variables, the direction of the effects is shown in parentheses. Bold type indicates the variables that were significant.

| Environment variables | adjusted R<sup>2</sup> | F | *p* | Spatial variables | adjusted R<sup>2</sup> | F | *p* |
| --- | --- | --- | --- | --- | --- | --- | --- |
| TP | <0.001 | 0.298 | 0.964 | Elevation (+) | **0.030** | **2.988** | **0.003** |
| TN | 0.004 | 0.645 | 0.743 | **MEM1 (-)** | **0.039** | **3.730** | **< 0.001** |
| Chl *a* | 0.011 | 1.309 | 0.218 | MEM2 | 0.018 | 1.940 | 0.066 |
| **WEF (PC1) (+)** | **0.069** | **6.367** | **< 0.001** | **MEM3 (+)** | **0.020** | **2.073** | **0.050** |
| **CYF (PC2) (-)** | **0.037** | **3.565** | **0.002** | **MEM4 (+)** | **0.028** | **2.767** | **0.006** |
| SDF (PC3) | 0.009 | 1.062 | 0.407 | MEM5 | 0.006 | 0.884 | 0.503 |
| **Watershed (+)** | **0.061** | **5.703** | **< 0.001** | MEM6 | 0.007 | 0.898 | 0.500 |
| **Volume (+)** | **0.059** | **5.504** | **< 0.001** | MEM7 | 0.016 | 1.721 | 0.099 |
| **Height (-)** | **0.021** | **2.158** | **0.035** | MEM8 | 0.009 | 1.101 | 0.343 |
| Width | 0.007 | 0.954 | 0.454 | **MEM9 (+)** | **0.043** | **4.132** | **0.001** |
| VWCF | 0.007 | 0.964 | 0.463 | **MEM10 (-)** | **0.051** | **4.841** | **< 0.001** |
| Turnover | 0.008 | 1.012 | 0.316 | MEM11 | 0.014 | 1.506 | 0.146 |
| **Age (+)** | **0.018** | **1.934** | **0.046** | MEM12 | 0.002 | 0.448 | 0.865 |

Also similar to the taxon-based structure, spatial variables for elevation and large- and small-scale MEMs significantly affected the trait-based community structure (Table 5). Among these variables, MEM 10 explained the most variation in community structure, followed by MEM 9 and MEM 1 (Table 5). Among the functional groups, large filter-feeding herbivores were positively associated with elevation and different scales of spatial configuration (MEM 1, 3, 4, 9, and 10). Similarly, large filter-feeding omnivores were positively associated with MEM 1, 9, and 10. In addition, small raptorial omnivores were positively associated with MEM 3 and 4, but negatively associated with MEM 1. Large browsing omnivores were negatively associated with MEM 4 (Table S9).

## Discussion

Our results show that when zooplankton communities were defined by traits, they were almost twice as well explained by environmental filters than by dispersal ones, whereas the contributions of these filters were indistinguishable for communities defined by taxonomy. These results support our hypothesis that environmental filters play a dominant role in determining functional community structure and that environmental and dispersal filters play relatively equal roles for taxonomic structure. Our results confirm that the relative importance of environmental and dispersal filters in determining community structure can vary with the descriptive unit of the community.

Among the variables related to the environmental filter, ordination axes characterizing fish communities had significant effects on the trait-based zooplankton communities. PC1, representing warm-water euryphagous fish assemblages (WEF) positively represented by largemouth bass and omnivorous cyprinids, significantly influenced both the taxon-based and trait-based community structures. Although these fishes often prey on zooplankton (Misawa, 2007), they are not purely planktivorous. As invasions of largemouth bass have resulted in decreased fish richness in Japanese lakes (Tsunoda & Mitsuo, 2012), it is likely that reservoirs with high values of WEF had fewer species of planktivorous fish, making predation pressure on zooplankton more moderate. In support of this, large zooplankton taxa that outcompete smaller zooplankton yet are favored by planktivorous fish (Brooks & Dodson, 1965; Hall et al. 1976; DeMott 1989; Urabe 1990) occurred in the trait-based community in such reservoirs. These larger zooplankton included large filter-feeding herbivores such as *Daphnia* and large omnivores such as calanoid copepods. The trait-based structure was also significantly affected by PC2, representing the cyprinid fish assemblage (CYF). Most of these cyprinid fish species are omnivorous but are known to aggressively feed on zooplankton (Sunaga, 1970; Urabe & Maruyama, 1986). Therefore, reservoirs with such fish communities are expected to select for zooplankton functional groups resistant to fish predation. Indeed, in reservoirs with high CYF scores, functional groups were dominant that can escape predation by quick swimming, such as the large filter-feeding omnivores. Calanoid copepods, the main components of this group, can sense vibrations from fish movement to expertly escape predation (Nassal et al., 1998). These results indicate that the top-down effects of fish strongly influence trait-based community structure.

Fish assemblages categorized as PC1 (WEF) and PC3 (SDF) also influenced the taxon-based community structure. However, the contribution of fish assemblages was much smaller for the taxon-based community than for the trait-based community. The classical size-efficiency hypothesis proposed that the size structure, rather than the species composition, of the zooplankton community is regulated by predators at higher trophic levels such as fish (Brooks and Dodson 1965). Since then, many studies have supported the adequacy of this hypothesis (e.g., Dodson 1974; Zaret 1980; Lynch 1979; H<u>ül</u>smann 2006; Ye et al. 2013). It follows that zooplankton communities structured by size class (i.e., traits) should be more affected by predators than those structured by taxonomy, but no study had yet demonstrated this. Importantly, our results clearly show that this is true for the zooplankton communities we studied. It should be noted that although the pond smelt *Hypomesus nipponiensis* is a plankton feeder (Yoshioka et al. 1994; Kawabata, 2002; Oh et al., 2019), the partial-dbRDA showed that the copepod *Cyclops vicinus* was positively related to SDF, while other *Cyclopidae* species were negatively related to SDF, although both were classified in the functional group of medium raptorial-feeding omnivores. These results suggest that the effects of fish predation are not necessarily the same among different species even within the same functional groups.

Among the local environmental variables, watershed area, water volume, and dam height also explained substantial portions of the variation in both community structures, but the effect of these variables differed among taxa and functional groups. For example, groups with large body sizes, such as large filter-feeding herbivores and omnivores, tended to occur in reservoirs with small watershed areas and those that were deep. As zooplankton can escape fish predation via vertical migration in deep reservoirs (Lampert, 1989; Gliwicz, 1986), they can likely harbor some functional groups with large body sizes. However, at the taxonomic level, although the large species *Daphnia galeata* was abundant in reservoirs with high dams (i.e. deep reservoir), other large zooplankton taxa, such as calanoid copepods, were rarely found in reservoirs with high dams. The results demonstrate clearly that although functional groups may have tight connections to certain environmental conditions, this may not be true for all species in that group.

According to Alfonso et al. (2010), species richness and abundance of copepods and cladocerans tended to increase up to 50 years after lake formation. In the present study, however, the trait-based community structure, rather than the taxon-based structure, was related with reservoir age. This suggests that functional diversity may have a stronger relationship with reservoir age than species richness.

Zooplankton can disperse passively by wind, flooding, and hitchhiking on mobile animals such as waterfowl (Vanschoenwinkel et al., 2008). Thus, it is likely that community structure is limited by the dispersal ability of zooplankton species. Indeed, the present study showed that the taxon-based community structure of zooplankton was substantially influenced by the spatial configuration of reservoirs at both smaller (MEM 9 to 11) and larger (MEM 2 to 5) scales. However, the effects of spatial variables varied among taxa. For example, some cladoceran species, such as *Bosminopsis deitersi*, *Daphnia galeata* and *D. dentifera*, showed high eigenvector values for one or more smaller spatial configurations, suggesting that they have limited dispersal abilities, or alternatively that they can easily disperse to neighboring reservoirs regardless of environmental conditions. On the other hand, some rotifers, such as *Ascomorpha ovalis*, *Cephalodella* sp. and *Collotheca* spp., showed high eigenvector values for the larger-scale spatial configurations (MEM 2 to 5), suggesting that they can disperse to more distant reservoirs. Although we lack direct knowledge about the dispersal abilities of these species, our results suggest that dispersal ability is highly variable among zooplankton taxa.

In a similar analysis, Padial et al. (2014) reported that environmental variables were more important than spatial variables in explaining variation in zooplankton communities defined by taxonomy among dozens of habitats in a floodplain. However, the present study that covers the Japanese archipelago is much greater in spatial scale and landscape diversity than a single floodplain, so our results are likely more reflective of general trends in zooplankton communities. In addition, our analysis may have underestimated the potential impact of dispersal filters by excluding less abundant species from the data. Nevertheless, the present study showed that dispersal and niche factors were equally important for zooplankton taxon-based community structures. Thus, the spatial configuration of habitats may be more important in determining the taxonomic structure of the zooplankton community.

Although they were less important compared to environmental variables, some Moran’s eigenvalues affected the trait-based community structure. For example, among the functional groups, large filter-feeding herbivores was strongly associated with both smaller (MEM 9 and 10) and larger spatial scales (MEM 1, 3, and 4), indicating that its distribution pattern is spatially complex, probably due to the inclusion of multiple species with different dispersal abilities in this group. However, some functional groups were associated with particular spatial scales: large and medium raptorial carnivores dispersed mainly at smaller spatial scales, whereas small and large filter-feeding omnivores dispersed mainly at larger spatial scales. Regardless of the fact that we did not consider dispersal ability when categorizing zooplankton taxa into functional groups, these results suggest that dispersal ability did indeed differ somewhat at a functional group level.

Although the rates of variation we were able to explain for zooplankton community structure using our variable sets were within previously described ranges for this system, our study does have some potential shortfalls. For example, Cottenie (2005) showed that multivariate analyses examining the effects of environmental and spatial variables explained 30 to 60% of the variation in animal and plant community structure in 95% of studies at the time. However, our analysis could not explain more than 50% of the total variation in taxon- and trait-based community structure. As sampling frequency varied among reservoirs, we defined our community structures based on annual averages. But as zooplankton community structure varies seasonally (Burns & Mitchell, 1980; Yoshida et al., 2001; Perera et al., 2021), the inclusion of seasonality may increase the resolution of the analysis when examining the relative importance between the effects of environmental and spatial variables on community structure. Another potential shortfall of our study is that we analyzed zooplankton communities using binary data, possibly causing us to miss some important factors affecting zooplankton structure. For example, we investigated the effects of vertical water circulation by aeration (VWCF) on the zooplankton community, which was installed in some reservoirs to reduce algal production. We initially suspected that VWCF may have affected some zooplankton species by physically disturbing them through water currents or movement, or by changing the composition and abundance of phytoplankton. However, we detected no significant effects of the VWCF on taxon- and trait-based zooplankton community structures, nor did we detect significant effects of TN, TP and Chl-*a* on these structures. However, it is often reported that taxon-based zooplankton community structures are related to the trophic status of the studied lakes through bottom-up effects (Pace, 1986; Yoshida et al., 2003; Perga et al., 2010). Lemmens et al. (2018) showed that in a shallow lake complex in Belgium, zooplankton biomass was mainly regulated by bottom-up effects, while zooplankton composition in terms of taxa was mainly determined by top-down effects from fish. Thus, the inclusion of continuous data may allow us to examine more deeply the effects of VWCR and trophic conditions on zooplankton community structure, and thus increase the relative importance of environmental variables in determining structures defined by taxonomy and function.

## Conclusion

Our extensive study of zooplankton in reservoirs showed that the relative importance of environmental and spatial filters in determining community structure varied depending on the unit used to describe the structure. Trait-based community structure was more influenced by environmental filters than by dispersal filters, but the taxonomic structure was equally regulated by both kinds of filters. These results suggest that even if the taxonomic structure of local zooplankton communities differs due to taxon-specific differences in range and dispersal ability, the communities tend to comprise functionally similar groups if the environmental conditions of local habitats are similar. However, because these results were obtained from binary data without considering the abundance of individual zooplankton taxa, the present study may have underestimated the effect of environmental filters in determining both taxon- and trait-based community structure compared to dispersal filters. Therefore, future studies seeking to tackle these important questions should examine whether the present finding is also supported when the abundance of composite organisms is considered.

## Acknowledgments

We thank the members of the Aquatic Ecology Laboratory at the Graduate School of Life Sciences, Tohoku University, for their help and comments. This study was financially supported by the Japan Society for the Promotion of Science (JSPS) Grant-in-Aid for Scientific Research (KAKENHI) 20H03315 and 23H02548, the Environment Research and Technology Development Fund (JPMEERF20214003) of the Environmental Restoration and Conservation Agency provided by the Ministry of the Environment of Japan, and a Dam-research fund by the Water Resources Environmental Center, Japan.

## Author Contributions

J. U. and H. S. planned and designed the study, H. S., J. U. and H. I. complied data, H. S. and J. U. performed analyses, H. S. and J. U. wrote the draft with J. M. K. providing major edits, and all authors contributed to the final manuscript.

