## Appendix S1 for "Differences in factors determining taxon-based and trait-based community structures: a field test using zooplankton"

Appendix S1
Supporting information for

### Differences in factors determining taxon-based and trait-based community structures: a field test using zooplankton

Hiromichi Suzuki, Hidetaka Ichiyanagi, Jamie M. Kass, and Jotaro Urabe

**Figure S1**
**Figure S2**
**Figure S3**

**Table S1**
**Table S2**
**Table S3**
**Table S4**
**Table S5**
**Table S6**
**Table S7**
**Table S8**
**Table S9**

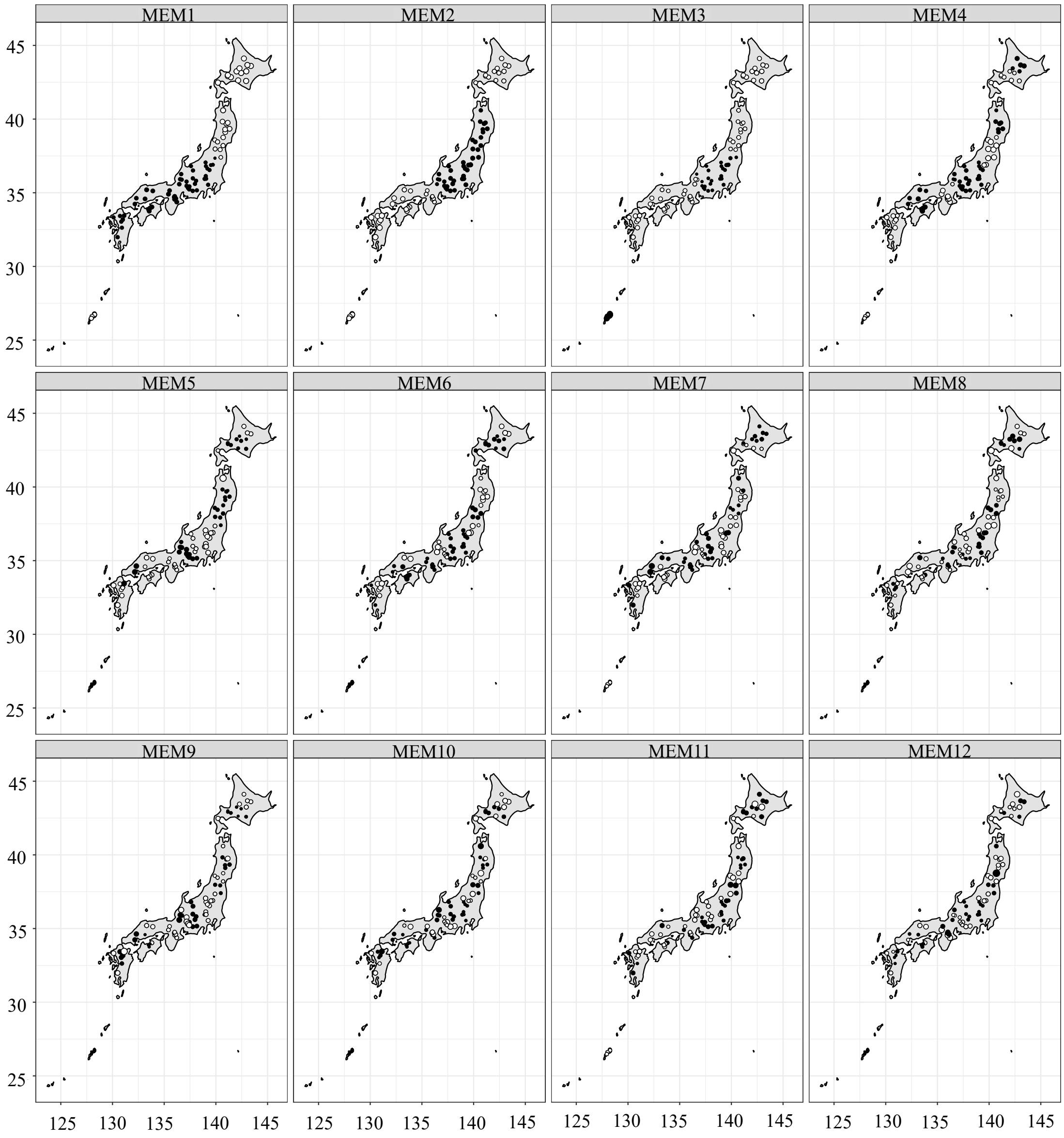


**Figure S1.** Maps showing Moran's eigenvalues from MEM 1 to MEM 12 for the reservoirs studied. The white and black circles represent positive and negative eigenvalues, respectively, and the size of the circle represents the magnitude of the absolute value.

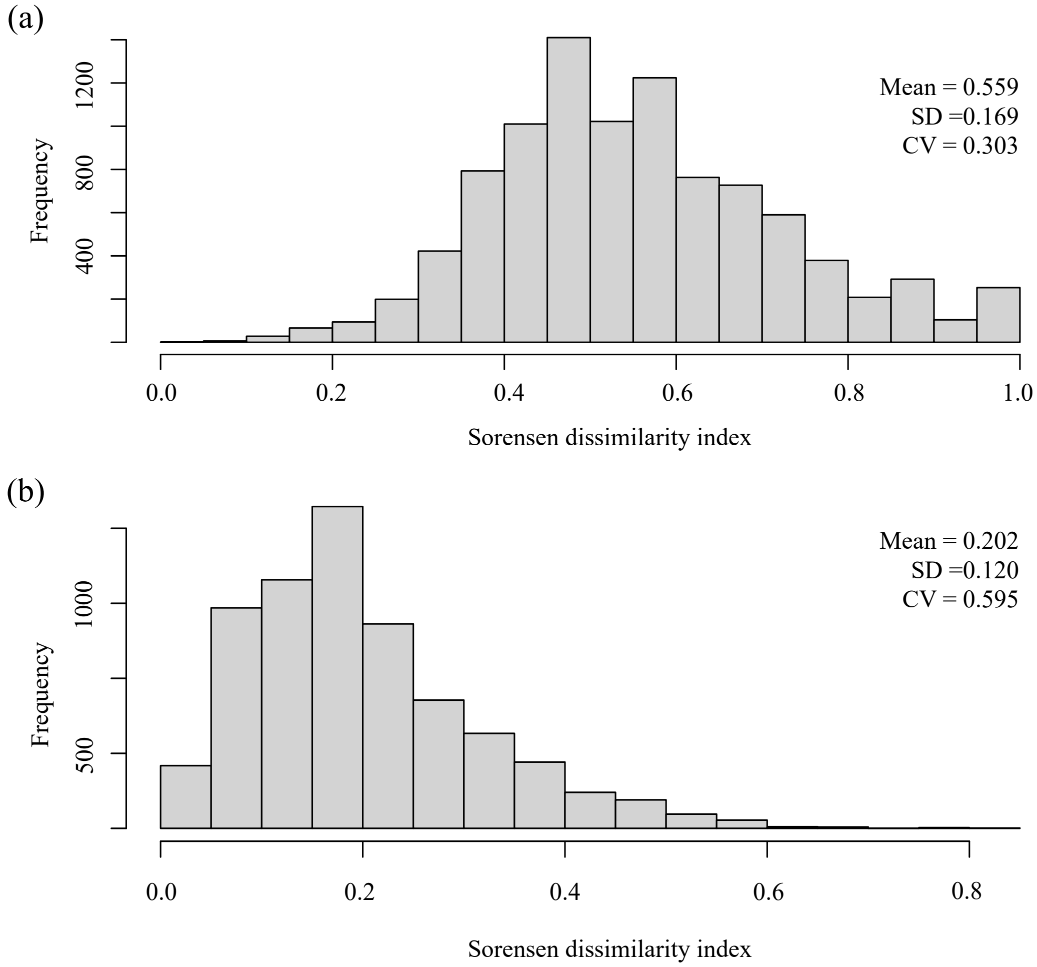


**Figure S2**. Histograms showing the frequencies of Sorensen dissimilarity index values among each of the taxa-based communities (a), and among each of the trait-based communities (b). Means, standard deviations (SD), and coefficients of variation (CV) of the index values are inserted in each panel.

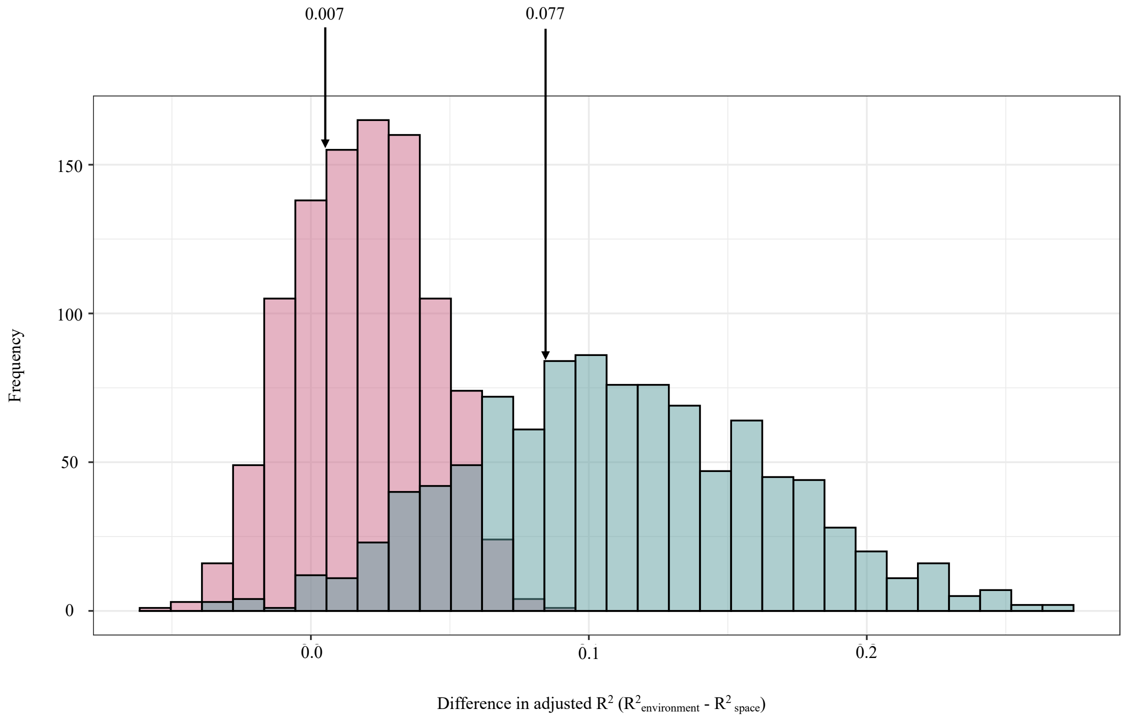


**Figure S3**. Histograms showing frequency distribution of difference in adjusted $R^2$ of environmental and spatial variables for taxon-based (red) and trait-based community structures (blue) estimated by randomization tests. Difference in adjusted $R^2$ estimated for the analysis of the dataset consisting of all data from all reservoirs is shown by the arrow.

**Table S1.** Moran’s eigenvalues (from MEM 1 to MEM 12) for each reservoir.

| Reservoirs | rounds | MEM1 | MEM2 | MEM3 | MEM4 | MEM5 | MEM6 | MEM7 | MEM8 | MEM9 | MEM10 | MEM11 | MEM12 |
|---|---|---|---|---|---|---|---|---|---|---|---|---|---|
| Agigawa | 3rd | -0.603 | -0.835 | -0.159 | -0.685 | -1.086 | -0.693 | -0.554 | -0.218 | -1.156 | 2.027 | -1.075 | 0.655 |
| Agigawa | 4th | -0.603 | -0.835 | -0.159 | -0.685 | -1.086 | -0.693 | -0.554 | -0.218 | -1.156 | 2.027 | -1.075 | 0.655 |
| Aha | 3rd | 0.539 | 1.137 | -3.333 | 0.139 | -0.206 | -0.193 | 0.088 | -0.059 | -0.054 | -0.077 | 0.099 | -0.064 |
| Arakawa | 3rd | 0.539 | 1.137 | -3.333 | 0.139 | -0.207 | -0.193 | 0.088 | -0.059 | -0.054 | -0.077 | 0.099 | -0.064 |
| Aseihigawa | 3rd | 1.489 | -0.600 | 0.254 | -0.211 | 3.835 | 0.173 | -1.151 | 1.395 | 0.123 | -3.186 | 1.732 | -0.880 |
| Aseihigawa | 4th | 1.489 | -0.600 | 0.254 | -0.211 | 3.835 | 0.173 | -1.151 | 1.395 | 0.123 | -3.186 | 1.732 | -0.880 |
| Benoki | 3rd | 0.539 | 1.137 | -3.333 | 0.139 | -0.206 | -0.193 | 0.088 | -0.059 | -0.054 | -0.077 | 0.099 | -0.064 |
| Fujiwara | 4th | -0.452 | -0.999 | -0.232 | -0.473 | 1.276 | -0.339 | 1.041 | -0.855 | 0.393 | -0.141 | -0.186 | -0.156 |
| Fukuchi | 3rd | 0.539 | 1.137 | -3.333 | 0.139 | -0.207 | -0.193 | 0.088 | -0.059 | -0.054 | -0.077 | 0.099 | -0.064 |
| Fukuchi | 4th | 0.539 | 1.137 | -3.333 | 0.139 | -0.207 | -0.193 | 0.088 | -0.059 | -0.054 | -0.077 | 0.099 | -0.064 |
| Fungawa | 3rd | 0.539 | 1.137 | -3.333 | 0.139 | -0.206 | -0.193 | 0.088 | -0.059 | -0.054 | -0.077 | 0.099 | -0.064 |
| Fungawa | 4th | 0.539 | 1.137 | -3.333 | 0.139 | -0.206 | -0.193 | 0.088 | -0.059 | -0.054 | -0.077 | 0.099 | -0.064 |
| Futase | 3rd | -0.488 | -0.946 | -0.183 | -0.596 | 0.820 | -0.483 | -0.107 | -0.071 | -0.381 | -0.422 | 0.213 | -0.238 |
| Futase | 4th | -0.488 | -0.946 | -0.183 | -0.596 | 0.820 | -0.483 | -0.107 | -0.071 | -0.381 | -0.422 | 0.213 | -0.238 |
| Gassan | 3rd | 0.612 | -1.043 | 0.355 | 1.838 | -0.635 | -1.293 | -0.210 | -1.320 | 0.076 | 0.057 | 1.826 | 0.941 |
| Gosyo | 3rd | 1.224 | -0.971 | 0.234 | -1.098 | -0.533 | 1.244 | 0.628 | 0.033 | -0.635 | -0.247 | -0.179 | 0.492 |
| Gosyo | 4th | 1.224 | -0.971 | 0.234 | -1.098 | -0.533 | 1.244 | 0.628 | 0.033 | -0.635 | -0.247 | -0.179 | 0.492 |
| Hachisu | 3rd | -1.035 | 0.233 | 0.333 | 1.210 | 0.211 | -0.373 | -0.286 | 0.104 | 0.235 | -0.297 | -0.380 | 2.054 |
| Hachisu | 4th | -1.035 | 0.233 | 0.333 | 1.210 | 0.211 | -0.373 | -0.286 | 0.104 | 0.235 | -0.297 | -0.380 | 2.054 |
| Hattabara | 3rd | -1.113 | 0.925 | 0.455 | -1.610 | 0.292 | -1.554 | 1.098 | 0.224 | -0.025 | -0.089 | 0.331 | -0.211 |
| Hanezi | 4th | 0.539 | 1.137 | -3.333 | 0.139 | -0.206 | -0.193 | 0.088 | -0.059 | -0.054 | -0.077 | 0.099 | -0.064 |
| Hinachi | 3rd | -1.013 | 0.199 | 0.362 | 1.251 | 0.124 | -0.276 | -0.384 | -0.257 | 0.261 | 0.872 | 0.131 | -1.509 |
| Hinachi | 4th | -1.013 | 0.199 | 0.362 | 1.251 | 0.124 | -0.276 | -0.384 | -0.257 | 0.261 | 0.872 | 0.131 | -1.509 |
| Hitokura | 3rd | -1.037 | 0.246 | 0.342 | 1.207 | 0.206 | -0.355 | -0.291 | 0.119 | 0.226 | -0.328 | -0.363 | 2.052 |
| Hitokura | 4th | -1.037 | 0.246 | 0.342 | 1.207 | 0.206 | -0.355 | -0.291 | 0.119 | 0.226 | -0.328 | -0.363 | 2.052 |
| Hiyoshi | 4th | -1.014 | 0.206 | 0.367 | 1.252 | 0.119 | -0.264 | -0.388 | -0.247 | 0.254 | 0.849 | 0.148 | -1.512 |
| Houheikyo | 4th | 1.816 | 0.514 | 0.809 | 0.412 | -0.441 | -1.000 | 0.050 | -0.197 | -0.188 | -0.765 | -1.255 | -0.363 |
| Ikari | 3rd | -0.202 | -1.272 | -0.522 | 1.114 | 0.686 | 2.469 | -1.530 | -0.837 | 0.203 | 0.063 | -0.448 | 0.231 |
| Ikeda | 3rd | -1.129 | 0.837 | 0.386 | -1.438 | 0.624 | -0.550 | 0.222 | -0.762 | 0.306 | -0.023 | -0.814 | 0.324 |
| Ikeda | 4th | -1.129 | 0.837 | 0.386 | -1.438 | 0.624 | -0.550 | 0.222 | -0.762 | 0.306 | -0.023 | -0.814 | 0.324 |
| Ishibuchi | 4th | 1.210 | -0.988 | 0.220 | -1.103 | -0.552 | 1.256 | 0.648 | 0.036 | -0.659 | -0.271 | -0.172 | 0.486 |
| Iwaonai | 4th | 1.670 | 0.882 | 0.615 | -1.553 | 0.597 | 1.181 | -0.418 | 0.741 | 0.324 | 0.088 | -1.461 | 2.415 |
| Iwaya | 3rd | -0.639 | -0.715 | -0.042 | -0.626 | -1.879 | 0.161 | 0.054 | 0.010 | 1.731 | 0.055 | -0.236 | 0.055 |
| Izarigawa | 3rd | 1.817 | 0.513 | 0.809 | 0.412 | -0.442 | -0.999 | 0.049 | -0.200 | -0.189 | -0.765 | -1.257 | -0.365 |
| Izarigawa | 4th | 1.817 | 0.513 | 0.809 | 0.412 | -0.442 | -0.999 | 0.049 | -0.200 | -0.189 | -0.765 | -1.257 | -0.365 |
| Jyozankei | 4th | 1.810 | 0.554 | 0.817 | 0.197 | -0.394 | -1.006 | -0.269 | -1.170 | -0.044 | -0.622 | -0.617 | 0.082 |
| Kamafusa | 3rd | 0.611 | -1.044 | 0.349 | 1.832 | -0.625 | -1.281 | -0.207 | -1.326 | 0.074 | 0.065 | 1.805 | 0.938 |
| Kamafusa | 4th | 0.611 | -1.044 | 0.349 | 1.832 | -0.625 | -1.281 | -0.207 | -1.326 | 0.074 | 0.065 | 1.805 | 0.938 |
| Kanayama | 3rd | 1.809 | 0.559 | 0.816 | 0.183 | -0.388 | -0.993 | -0.285 | -1.178 | -0.034 | -0.592 | -0.623 | 0.084 |
| Kanayama | 4th | 1.809 | 0.559 | 0.816 | 0.183 | -0.388 | -0.993 | -0.285 | -1.178 | -0.034 | -0.592 | -0.623 | 0.084 |
| Kanna | 3rd | 0.539 | 1.137 | -3.333 | 0.139 | -0.206 | -0.193 | 0.088 | -0.059 | -0.054 | -0.077 | 0.099 | -0.064 |
| Kanna | 4th | 0.539 | 1.137 | -3.333 | 0.139 | -0.206 | -0.193 | 0.088 | -0.059 | -0.054 | -0.077 | 0.099 | -0.064 |
| Kanoko | 3rd | 1.694 | 0.849 | 0.645 | -1.523 | 0.511 | 1.292 | -0.513 | 0.378 | 0.344 | 1.260 | -0.955 | -1.169 |
| Kanoko | 4th | 1.694 | 0.849 | 0.645 | -1.523 | 0.511 | 1.292 | -0.513 | 0.378 | 0.344 | 1.260 | -0.955 | -1.169 |
| Katsurasawa | 3rd | 1.809 | 0.559 | 0.816 | 0.185 | -0.388 | -0.995 | -0.283 | -1.174 | -0.033 | -0.593 | -0.623 | 0.087 |
| Katsurasawa | 4th | 1.809 | 0.559 | 0.816 | 0.185 | -0.388 | -0.995 | -0.283 | -1.174 | -0.033 | -0.593 | -0.623 | 0.087 |
| Kawaji | 3rd | -0.202 | -1.272 | -0.522 | 1.114 | 0.686 | 2.468 | -1.530 | -0.837 | 0.203 | 0.063 | -0.449 | 0.231 |
| Kawaji | 4th | -0.202 | -1.272 | -0.522 | 1.114 | 0.686 | 2.468 | -1.530 | -0.837 | 0.203 | 0.063 | -0.449 | 0.231 |
| Kawamata | 3rd | -0.258 | -1.216 | -0.465 | 0.792 | 0.909 | 2.213 | -1.396 | -1.379 | 0.436 | 0.051 | -1.214 | 0.550 |
| Kawamata | 4th | -0.258 | -1.216 | -0.465 | 0.792 | 0.909 | 2.213 | -1.396 | -1.379 | 0.436 | 0.051 | -1.214 | 0.550 |
| Koshibu | 3rd | -0.525 | -0.885 | -0.127 | -0.717 | 0.303 | -0.581 | -1.261 | 0.756 | -1.066 | -0.589 | 0.540 | -0.205 |
| Koshibu | 4th | -0.525 | -0.885 | -0.127 | -0.717 | 0.303 | -0.581 | -1.261 | 0.756 | -1.066 | -0.589 | 0.540 | -0.205 |
| Kusaki | 3rd | -0.451 | -1.000 | -0.235 | -0.469 | 1.273 | -0.339 | 1.039 | -0.856 | 0.393 | -0.130 | -0.194 | -0.156 |
| Kusaki | 4th | -0.451 | -1.000 | -0.235 | -0.469 | 1.273 | -0.339 | 1.039 | -0.856 | 0.393 | -0.130 | -0.194 | -0.156 |
| Kuzuryu | 3rd | -0.700 | -0.439 | 0.201 | -0.225 | -1.617 | 1.518 | 1.669 | -0.555 | -1.264 | -0.692 | 0.443 | 0.049 |
| Kyuuragi | 3rd | -0.533 | 2.068 | 0.740 | 0.915 | 1.410 | -0.142 | -1.250 | 0.611 | 1.117 | 0.887 | -1.022 | 0.387 |
| Managawa | 3rd | -0.716 | -0.368 | 0.261 | -0.122 | -1.498 | 1.777 | 1.906 | -0.626 | -1.838 | -0.766 | 0.517 | 0.037 |
| Maruyama | 3rd | -0.641 | -0.711 | -0.041 | -0.629 | -1.879 | 0.149 | 0.048 | 0.012 | 1.744 | 0.056 | -0.236 | 0.051 |
| Maruyama | 4th | -0.641 | -0.711 | -0.041 | -0.629 | -1.879 | 0.149 | 0.048 | 0.012 | 1.744 | 0.056 | -0.236 | 0.051 |
| Matsubara | 3rd | -0.612 | 2.029 | 0.826 | 0.914 | 0.376 | 1.463 | 1.279 | -0.409 | -2.019 | -0.442 | 0.356 | -0.002 |
| Matsubara | 4th | -0.612 | 2.029 | 0.826 | 0.914 | 0.376 | 1.463 | 1.279 | -0.409 | -2.019 | -0.442 | 0.356 | -0.002 |
| Midorikawa | 4th | -0.574 | 2.053 | 0.788 | 0.949 | 0.942 | 0.695 | 0.142 | 0.017 | -0.849 | 0.153 | -0.098 | -0.040 |
| Miharu | 3rd | 0.244 | -1.213 | -0.125 | 1.900 | -0.254 | 0.033 | 1.003 | 2.768 | -0.382 | -0.341 | -1.370 | -0.911 |
| Misogawa | 3rd | -0.523 | -0.887 | -0.126 | -0.718 | 0.307 | -0.570 | -1.264 | 0.765 | -1.072 | -0.613 | 0.556 | -0.208 |
| Misogawa | 4th | -0.523 | -0.887 | -0.126 | -0.718 | 0.307 | -0.570 | -1.264 | 0.765 | -1.072 | -0.613 | 0.556 | -0.208 |
| Miwa | 3rd | -0.522 | -0.889 | -0.128 | -0.712 | 0.310 | -0.566 | -1.255 | 0.758 | -1.068 | -0.606 | 0.549 | -0.208 |
| Miwa | 4th | -0.522 | -0.889 | -0.128 | -0.712 | 0.310 | -0.566 | -1.255 | 0.758 | -1.068 | -0.606 | 0.549 | -0.208 |
| Miyagase | 3rd | -0.464 | -0.987 | -0.232 | -0.482 | 1.258 | -0.396 | 1.045 | -0.886 | 0.418 | -0.050 | -0.241 | -0.143 |
| Miyagase | 4th | -0.464 | -0.987 | -0.232 | -0.482 | 1.258 | -0.396 | 1.045 | -0.886 | 0.418 | -0.050 | -0.241 | -0.143 |

**Table S1** continued

| Reservoirs | rounds | MEM1 | MEM2 | MEM3 | MEM4 | MEM5 | MEM6 | MEM7 | MEM8 | MEM9 | MEM10 | MEM11 | MEM12 |
|---|---|---|---|---|---|---|---|---|---|---|---|---|---|
| Muroo | 3rd | -1.036 | 0.236 | 0.335 | 1.211 | 0.209 | -0.369 | -0.288 | 0.108 | 0.232 | -0.306 | -0.374 | 2.055 |
| Muroo | 4th | -1.036 | 0.236 | 0.335 | 1.211 | 0.209 | -0.369 | -0.288 | 0.108 | 0.232 | -0.306 | -0.374 | 2.055 |
| Nagashima | 4th | -0.598 | -0.840 | -0.165 | -0.670 | -1.077 | -0.676 | -0.532 | -0.230 | -1.148 | 2.035 | -1.086 | 0.652 |
| Narugo | 4th | 0.940 | -0.969 | 0.400 | 0.329 | -0.474 | 0.416 | -0.306 | -1.027 | 0.075 | 2.873 | 1.081 | -7.320 |
| Nibutani | 3rd | 1.818 | 0.428 | 0.752 | 0.769 | -0.322 | -0.237 | 0.461 | 2.164 | -0.012 | 0.777 | 0.224 | 0.196 |
| Nukui | 4th | -0.976 | 1.332 | 0.643 | -0.987 | -2.462 | -0.401 | -3.635 | 2.306 | -2.142 | -0.852 | 0.942 | -0.191 |
| Nunome | 3rd | -1.014 | 0.201 | 0.363 | 1.251 | 0.123 | -0.273 | -0.385 | -0.254 | 0.260 | 0.867 | 0.134 | -1.509 |
| Nunome | 4th | -1.014 | 0.201 | 0.363 | 1.251 | 0.123 | -0.273 | -0.385 | -0.254 | 0.260 | 0.867 | 0.134 | -1.509 |
| Ookawa | 3rd | -0.009 | -1.280 | -0.401 | 1.597 | 0.337 | 1.635 | 0.653 | 3.369 | 0.241 | 1.885 | 1.589 | 0.582 |
| Ookawa | 4th | -0.009 | -1.280 | -0.401 | 1.597 | 0.337 | 1.635 | 0.653 | 3.369 | 0.241 | 1.885 | 1.589 | 0.582 |
| Oomachi | 3rd | -0.516 | -0.894 | -0.127 | -0.711 | 0.319 | -0.540 | -1.257 | 0.773 | -1.079 | -0.652 | 0.577 | -0.214 |
| Oomachi | 4th | -0.516 | -0.894 | -0.127 | -0.711 | 0.319 | -0.540 | -1.257 | 0.773 | -1.079 | -0.652 | 0.577 | -0.214 |
| Origawa | 3rd | -0.641 | -0.711 | -0.042 | -0.628 | -1.875 | 0.146 | 0.049 | 0.009 | 1.743 | 0.062 | -0.240 | 0.051 |
| Origawa | 4th | -0.641 | -0.711 | -0.042 | -0.628 | -1.875 | 0.146 | 0.049 | 0.009 | 1.743 | 0.062 | -0.240 | 0.051 |
| Pirika | 3rd | 1.798 | 0.356 | 0.730 | 0.723 | 0.156 | -0.506 | 1.927 | 3.204 | 0.595 | 1.897 | 0.621 | 0.922 |
| Pirika | 4th | 1.798 | 0.356 | 0.730 | 0.723 | 0.156 | -0.506 | 1.927 | 3.204 | 0.595 | 1.897 | 0.621 | 0.922 |
| Ryuumon | 4th | -0.593 | 2.042 | 0.810 | 0.936 | 0.670 | 1.110 | 0.746 | -0.207 | -1.540 | -0.160 | 0.146 | -0.023 |
| Sagae | 3rd | 0.612 | -1.044 | 0.353 | 1.837 | -0.632 | -1.290 | -0.209 | -1.322 | 0.075 | 0.060 | 1.819 | 0.940 |
| Sagae | 4th | 0.612 | -1.044 | 0.353 | 1.837 | -0.632 | -1.290 | -0.209 | -1.322 | 0.075 | 0.060 | 1.819 | 0.940 |
| Sagurigawa | 3rd | -0.362 | -0.977 | -0.128 | -0.309 | 1.335 | -0.562 | 2.263 | 1.408 | 0.831 | 1.396 | 0.404 | 0.741 |
| Sagurigawa | 4th | -0.362 | -0.977 | -0.128 | -0.309 | 1.335 | -0.562 | 2.263 | 1.408 | 0.831 | 1.396 | 0.404 | 0.741 |
| Sameura | 3rd | -1.112 | 0.927 | 0.456 | -1.604 | 0.293 | -1.560 | 1.098 | 0.218 | -0.020 | -0.069 | 0.314 | -0.208 |
| Sameura | 4th | -1.112 | 0.927 | 0.456 | -1.604 | 0.293 | -1.560 | 1.098 | 0.218 | -0.020 | -0.069 | 0.314 | -0.208 |
| Satsunaigawa | 3rd | 1.816 | 0.514 | 0.808 | 0.405 | -0.442 | -0.989 | 0.039 | -0.214 | -0.188 | -0.754 | -1.259 | -0.371 |
| Satsunaigawa | 4th | 1.816 | 0.514 | 0.808 | 0.405 | -0.442 | -0.989 | 0.039 | -0.214 | -0.188 | -0.754 | -1.259 | -0.371 |
| Shijushida | 4th | 1.263 | -0.950 | 0.192 | -1.121 | -0.053 | 0.388 | -0.767 | 0.619 | 1.343 | 0.510 | -1.109 | 0.913 |
| Shimokubo | 3rd | -0.461 | -0.993 | -0.231 | -0.481 | 1.267 | -0.376 | 1.045 | -0.875 | 0.409 | -0.090 | -0.217 | -0.148 |
| Shimokubo | 4th | -0.461 | -0.993 | -0.231 | -0.481 | 1.267 | -0.376 | 1.045 | -0.875 | 0.409 | -0.090 | -0.217 | -0.148 |
| Shiouke | 3rd | -0.612 | 2.029 | 0.826 | 0.914 | 0.377 | 1.463 | 1.279 | -0.408 | -2.017 | -0.442 | 0.356 | -0.002 |
| Shinguu | 3rd | -1.114 | 0.923 | 0.455 | -1.610 | 0.294 | -1.569 | 1.103 | 0.215 | -0.020 | -0.069 | 0.319 | -0.211 |
| Shinguu | 4th | -1.114 | 0.923 | 0.455 | -1.610 | 0.294 | -1.569 | 1.103 | 0.215 | -0.020 | -0.069 | 0.319 | -0.211 |
| Shintoyone | 3rd | -0.602 | -0.834 | -0.162 | -0.680 | -1.081 | -0.695 | -0.548 | -0.223 | -1.149 | 2.034 | -1.081 | 0.653 |
| Shintoyone | 4th | -0.602 | -0.834 | -0.162 | -0.680 | -1.081 | -0.695 | -0.548 | -0.223 | -1.149 | 2.034 | -1.081 | 0.653 |
| Shirakawa | 3rd | 0.468 | -1.119 | 0.177 | 2.009 | -0.607 | -1.134 | 0.572 | 0.737 | -0.452 | -1.595 | -2.822 | -0.708 |
| Shirakawa | 4th | 0.468 | -1.119 | 0.177 | 2.009 | -0.607 | -1.134 | 0.572 | 0.737 | -0.452 | -1.595 | -2.822 | -0.708 |
| Sonohara | 4th | -0.453 | -0.999 | -0.233 | -0.472 | 1.274 | -0.342 | 1.041 | -0.857 | 0.394 | -0.132 | -0.192 | -0.155 |
| Sugesawa | 3rd | -1.141 | 0.706 | 0.268 | -1.070 | 0.996 | 1.186 | -1.242 | -1.643 | 0.529 | 0.029 | -1.425 | 0.661 |
| Sugesawa | 4th | -1.141 | 0.706 | 0.268 | -1.070 | 0.996 | 1.186 | -1.242 | -1.643 | 0.529 | 0.029 | -1.425 | 0.661 |
| Surikamigawa | 4th | 0.472 | -1.117 | 0.176 | 2.004 | -0.600 | -1.133 | 0.578 | 0.735 | -0.455 | -1.604 | -2.833 | -0.705 |
| Syorenji | 3rd | -1.013 | 0.199 | 0.362 | 1.251 | 0.124 | -0.277 | -0.384 | -0.257 | 0.261 | 0.873 | 0.131 | -1.509 |
| Syorenji | 4th | -1.013 | 0.199 | 0.362 | 1.251 | 0.124 | -0.277 | -0.384 | -0.257 | 0.261 | 0.873 | 0.131 | -1.509 |
| Taisetsu | 3rd | 1.695 | 0.849 | 0.646 | -1.523 | 0.512 | 1.291 | -0.511 | 0.382 | 0.344 | 1.259 | -0.954 | -1.169 |
| Taisetsu | 4th | 1.695 | 0.849 | 0.646 | -1.523 | 0.512 | 1.291 | -0.511 | 0.382 | 0.344 | 1.259 | -0.954 | -1.169 |
| Takayama | 3rd | -1.013 | 0.199 | 0.363 | 1.253 | 0.123 | -0.275 | -0.385 | -0.256 | 0.260 | 0.868 | 0.135 | -1.512 |
| Takayama | 4th | -1.013 | 0.199 | 0.363 | 1.253 | 0.123 | -0.275 | -0.385 | -0.256 | 0.260 | 0.868 | 0.135 | -1.512 |
| Takisato | 4th | 1.773 | 0.673 | 0.776 | -0.494 | -0.057 | -0.202 | -0.834 | -1.946 | 0.478 | 1.378 | 3.310 | 1.119 |
| Tamagawa | 4th | 1.224 | -0.971 | 0.236 | -1.100 | -0.538 | 1.242 | 0.629 | 0.038 | -0.637 | -0.253 | -0.172 | 0.493 |
| Tase | 3rd | 1.218 | -0.977 | 0.228 | -1.098 | -0.534 | 1.249 | 0.634 | 0.029 | -0.643 | -0.250 | -0.182 | 0.488 |
| Tase | 4th | 1.218 | -0.977 | 0.228 | -1.098 | -0.534 | 1.249 | 0.634 | 0.029 | -0.643 | -0.250 | -0.182 | 0.488 |
| Tedorigawa | 3rd | -0.566 | -0.752 | -0.003 | -0.678 | -0.500 | 0.251 | -0.684 | 1.005 | 1.825 | -2.587 | 1.401 | -0.806 |
| Tedorigawa | 4th | -0.566 | -0.752 | -0.003 | -0.678 | -0.500 | 0.251 | -0.684 | 1.005 | 1.825 | -2.587 | 1.401 | -0.806 |
| Teraushi | 4th | -0.611 | 2.028 | 0.825 | 0.911 | 0.376 | 1.466 | 1.275 | -0.406 | -2.017 | -0.446 | 0.355 | 0.000 |
| Tokachi | 3rd | 1.773 | 0.672 | 0.776 | -0.495 | -0.059 | -0.199 | -0.837 | -1.953 | 0.476 | 1.378 | 3.312 | 1.115 |
| Tokachi | 4th | 1.773 | 0.672 | 0.776 | -0.495 | -0.059 | -0.199 | -0.837 | -1.953 | 0.476 | 1.378 | 3.312 | 1.115 |
| Tomada | 4th | -1.146 | 0.608 | 0.204 | -0.719 | 1.130 | 1.944 | -0.833 | 0.707 | 0.719 | 1.600 | 0.048 | 1.233 |
| Tomisato | 3rd | -1.113 | 0.927 | 0.456 | -1.605 | 0.293 | -1.559 | 1.098 | 0.219 | -0.021 | -0.073 | 0.317 | -0.208 |
| Tomisato | 4th | -1.113 | 0.927 | 0.456 | -1.605 | 0.293 | -1.559 | 1.098 | 0.219 | -0.021 | -0.073 | 0.317 | -0.208 |
| Tsuruta | 3rd | -0.529 | 2.064 | 0.734 | 0.913 | 1.410 | -0.141 | -1.229 | 0.613 | 1.113 | 0.881 | -1.019 | 0.385 |
| Tsuruta | 4th | -0.529 | 2.064 | 0.734 | 0.913 | 1.410 | -0.141 | -1.229 | 0.613 | 1.113 | 0.881 | -1.019 | 0.385 |
| Unaduki | 3rd | -0.514 | -0.893 | -0.127 | -0.711 | 0.321 | -0.532 | -1.260 | 0.779 | -1.083 | -0.670 | 0.588 | -0.215 |
| Urayama | 3rd | -0.462 | -0.991 | -0.231 | -0.482 | 1.265 | -0.384 | 1.045 | -0.879 | 0.413 | -0.077 | -0.225 | -0.146 |
| Urayama | 4th | -0.462 | -0.991 | -0.231 | -0.482 | 1.265 | -0.384 | 1.045 | -0.879 | 0.413 | -0.077 | -0.225 | -0.146 |
| Yabakei | 3rd | -0.750 | 1.814 | 0.807 | 0.271 | -2.182 | 1.825 | 0.959 | 0.318 | 3.882 | -2.395 | 0.891 | -0.617 |
| Yabakei | 4th | -0.750 | 1.814 | 0.807 | 0.271 | -2.182 | 1.825 | 0.959 | 0.318 | 3.882 | -2.395 | 0.891 | -0.617 |
| Yahagi | 3rd | -0.641 | -0.711 | -0.043 | -0.626 | -1.872 | 0.145 | 0.050 | 0.007 | 1.741 | 0.067 | -0.243 | 0.051 |
| Yahagi | 4th | -0.641 | -0.711 | -0.043 | -0.626 | -1.872 | 0.145 | 0.050 | 0.007 | 1.741 | 0.067 | -0.243 | 0.051 |
| Yasaka | 4th | -0.951 | 1.400 | 0.671 | -0.847 | -2.741 | -0.169 | -3.546 | 2.349 | -1.846 | -0.503 | 0.733 | 0.046 |
| Yokoyama | 3rd | -0.764 | -0.260 | 0.307 | 0.107 | -1.318 | 1.840 | 2.194 | -0.837 | -2.653 | -0.919 | 0.994 | -0.432 |
| Yokoyama | 4th | -0.764 | -0.260 | 0.307 | 0.107 | -1.318 | 1.840 | 2.194 | -0.837 | -2.653 | -0.919 | 0.994 | -0.432 |
| Yuda | 4th | 1.214 | -0.984 | 0.224 | -1.102 | -0.547 | 1.252 | 0.642 | 0.036 | -0.652 | -0.265 | -0.173 | 0.488 |

**Table S2.** Zooplankton taxa and their functional groups. Taxa appearing in more than 15% of the lists are marked with an asterisk.

| Taxa | Body size | Feeding modes | Food habits | Population growth rate | Functional group |
|---|---|---|---|---|---|
| *Alona* spp.* | Small | Filter feeder | Detriti- / Herbivore | middle | SFDH |
| *A. rectangula* | Small | Filter feeder | Detriti- / Herbivore | middle | SFDH |
| *A.* gutata | Small | Filter feeder | Detriti- / Herbivore | middle | SFDH |
| *Bosmina coregoni* | Small | Filter feeder | Herbivore | middle | SFDH |
| *B. fatalis* | Small | Filter feeder | Herbivore | middle | SFDH |
| *B. longilostris** | Small | Filter feeder | Herbivore | middle | SFDH |
| *Bosminopsis deiteris** | Small | Filter feeder | Herbivore | middle | SFDH |
| *Camptocercus rectirostris* | Middle | Filter feeder | Herbivore | middle | MFH |
| *Ceriodaphnia* sp.* | Middle | Filter feeder | Herbivore | middle | MFH |
| *C. quadrangula* | Middle | Filter feeder | Herbivore | middle | MFH |
| *C. pulchella* | Middle | Filter feeder | Herbivore | middle | MFH |
| *Chydorus* sp. | Small | Filter feeder | Detriti- / Herbivore | middle | SFDH |
| *C. sphaericus** | Small | Filter feeder | Detriti- / Herbivore | middle | SFDH |
| *Daphnia* sp. | Large | Filter feeder | Herbivore | middle | LFH |
| *D. galeata** | Large | Filter feeder | Herbivore | middle | LFH |
| *D. dentifera** | Large | Filter feeder | Herbivore | middle | LFH |
| *D. pulex* | Large | Filter feeder | Herbivore | middle | LFH |
| *D. ambigua* | Large | Filter feeder | Herbivore | middle | LFH |
| *Diaphanosoma* sp. | Middle | Filter feeder | Herbivore | middle | MFH |
| *D. brachyurum** | Middle | Filter feeder | Herbivore | middle | MFH |
| *Holopedium gibberum* | Large | Filter feeder | Herbivore | middle | LFH |
| *Ilyocryptus sordidus* | Small | Filter feeder | Detriti- / Herbivore | middle | SFDH |
| *Leptodora kinditii* | Large | Raptorial feeder | Carnivore | middle | LRC |
| *Leydigia ciliate* | Small | Filter feeder | Detriti- / Herbivore | middle | SFD |
| *L. leydigii* | Small | Filter feeder | Detriti- / Herbivore | middle | SFD |
| *Moina* sp. | Middle | Filter feeder | Herbivore | middle | MFH |
| *M. macrocopa* | Middle | Filter feeder | Herbivore | middle | MFH |
| *Monospilus dispar* | Small | Filter feeder | Detriti- / Herbivore | middle | SFDH |
| *Pleuroxus* sp. | Small | Filter feeder | Detriti- / Herbivore | middle | SFDH |
| *Polyphemus pediculus* | Large | Raptorial feeder | Carnivore | middle | LRC |
| Scapholeberis sp. | Middle | Filter feeder | Herbivore | middle | MFD |
| *Sida crystalina* | Large | Filter feeder | Herbivore | middle | LFH |
| *Simocephalus* sp. | Large | Filter feeder | Herbivore | middle | LFH |

**Table S2** continued

| Taxa | Body size | Feeding modes | Food habits | Population growth rate | Functional group |
|---|---|---|---|---|---|
| *Acathocyclops* sp. | Middle | Raptorial feeder | Omnivore | low | MRO |
| *Acanthodiaptomus pacificus* | Large | Filter feeder | Omnivore | low | LFO |
| *Bryocamptus* sp. | Large | Browser | Omnivore | low | LBO |
| *Calanoida** | Large | Filter feeder | Omnivore | low | LFO |
| *Canthocamptus* sp. | Large | Browser | Omnivore | low | LBO |
| *Diaptomidae* | Large | Filter feeder | Omnivore | low | LFO |
| *Cyclopidae** | Middle | Raptorial feeder | Omnivore | low | MRO |
| *Harpacticoida* | Large | Browser | Omnivore | low | LBO |
| *Cyclops kikuchii* | Middle | Raptorial feeder | Omnivore | low | MRO |
| *C. strenuus* | Middle | Raptorial feeder | Omnivore | low | MRO |
| *C. vicinus** | Middle | Raptorial feeder | Omnivore | low | MRO |
| *Diacyclops* sp. | Middle | Raptorial feeder | Omnivore | low | MRO |
| *D. bicuspidatus* | Middle | Raptorial feeder | Omnivore | low | MRO |
| *Eodiaptomus japonicus** | Large | Filter feeder | Omnivore | low | LFO |
| *Eucyclops* sp. | Middle | Raptorial feeder | Omnivore | low | MRO |
| *E. serrulatus* | Middle | Raptorial feeder | Omnivore | low | MRO |
| *Eurytemora affinis* | Large | Filter feeder | Omnivore | low | LFO |
| *Limnoncaea* sp. | Middle | Raptorial feeder | Omnivore | low | MRO |
| *Megacyclops* sp. | Middle | Raptorial feeder | Omnivore | low | MRO |
| *Mesocyclops* sp. | Middle | Raptorial feeder | Omnivore | low | MRO |
| *M. leuckarti* | Middle | Raptorial feeder | Omnivore | low | MRO |
| *Microcyclops varicans* | Middle | Raptorial feeder | Omnivore | low | MRO |
| *Moraria* sp. | Large | Browser | Omnivore | low | LBO |
| *Paracyclops fimbriatus* | Middle | Raptorial feeder | Omnivore | low | MRO |
| *Thermocyclops* sp. | Middle | Raptorial feeder | Omnivore | low | MRO |
| *T. crassus* | Middle | Raptorial feeder | Omnivore | low | MRO |
| *T. hyalinus* | Middle | Raptorial feeder | Omnivore | low | MRO |
| *T. taihokuensis* | Middle | Raptorial feeder | Omnivore | low | MRO |
| *Tropocyclops* sp. | Middle | Raptorial feeder | Omnivore | low | MRO |
| *Auraeopsis* sp. | Small | Filter feeder | Detritivore | high | SFD |
| *Ascomorpha* spp.* | Small | Raptorial feeder | Herbivore | high | SRH |
| *A. ovalis** | Small | Raptorial feeder | Herbivore | high | SRH |
| *Asplanchna* sp.* | Middle | Raptorial feeder | Carnivore | high | MRC |
| *A. priodonta** | Middle | Raptorial feeder | Carnivore | high | MRC |

**Table S2** continued

| Taxa | Body size | Feeding modes | Food habits | Population growth rate | Functional group |
|---|---|---|---|---|---|
| *Bdelloidae* | Small | Filter feeder | Detritivore | high | SFD |
| *Brachionus* sp. | Small | Filter feeder | Detritivore | high | SFD |
| *B. calciflorus* | Small | Filter feeder | Detritivore | high | SFD |
| *B. quadridentatus* | Small | Filter feeder | Detritivore | high | SFD |
| *Cephalodella* sp.* | Small | Raptorial feeder | Omnivore | high | SRO |
| *Collotheca* spp.* | Small | Filter feeder | Detritivore | high | SFD |
| *Colurella* sp.* | Small | Filter feeder | Detritivore | high | SFD |
| *Conochiloides* spp.* | Small | Filter feeder | Detritivore | high | SFD |
| *Conochilus* spp.* | Small | Filter feeder | Detritivore | high | SFD |
| *Dipleuchlanis propatula* | Small | Filter feeder | Detritivore | high | SFD |
| *Euchlanis* sp. | Small | Filter feeder | Detritivore | high | SFD |
| *E. dilatate** | Small | Filter feeder | Detritivore | high | SFD |
| *Filinia* sp. | Small | Filter feeder | Detritivore | high | SFD |
| *F. longiseta** | Small | Filter feeder | Detritivore | high | SFD |
| *Gastropus* sp. | Small | Raptorial feeder | Herbivore | high | SRH |
| *Habrotrocha* sp. | Small | Filter feeder | Detritivore | high | SFD |
| *Hexarthra mira** | Small | Filter feeder | Detritivore | high | SFD |
| *Kellicottia bostoniensis* | Small | Filter feeder | Detritivore | high | SFD |
| *K. longispina** | Small | Filter feeder | Detritivore | high | SFD |
| *Keratella sp.* | Small | Filter feeder | Detritivore | high | SFD |
| *K. cochlearis** | Small | Filter feeder | Detritivore | high | SFD |
| *K. quadrata** | Small | Filter feeder | Detritivore | high | SFD |
| *K. valga* | Small | Filter feeder | Detritivore | high | SFD |
| *Lecane* sp. | Small | Filter feeder | Detritivore | high | SFD |
| *L. luna* | Small | Filter feeder | Detritivore | high | SFD |
| *Lepadella* sp. | Small | Filter feeder | Detritivore | high | SFD |
| *Macrochaetus subquadratus* | Small | Filter feeder | Detritivore | high | SFD |
| *Monommata* sp. | Small | Raptorial feeder | Herbivore | high | SRH |

**Table S2** continued

| Taxa | Body size | Feeding modes | Food habits | Population growth rate | Functional group |
|---|---|---|---|---|---|
| *Monostyla* sp. | Small | Filter feeder | Detritivore | high | SFD |
| *Mytilina* sp. | Small | Filter feeder | Detritivore | high | SFD |
| *Notholca* sp. | Small | Filter feeder | Herbivore | high | SFD |
| *N. labis* | Small | Filter feeder | Herbivore | high | SFD |
| *Notommata* sp. | Small | Raptorial feeder | Omnivore | high | SRO |
| *Notommatidae* | Small | Raptorial feeder | Omnivore | high | SRO |
| *Philodinidae** | Small | Filter feeder | Detritivore | high | SFD |
| *Platyias* sp. | Small | Filter feeder | Detritivore | high | SFD |
| *P.* patulus | Small | Filter feeder | Detritivore | high | SFD |
| *Ploesoma sp.* | Small | Raptorial feeder | Omnivore | high | SRO |
| *P. hudsoni** | Small | Raptorial feeder | Omnivore | high | SRO |
| *P. truncatum** | Small | Raptorial feeder | Omnivore | high | SRO |
| *Polyarthra sp.* | Small | Raptorial feeder | Herbivore | high | SRH |
| *P. euryptera** | Small | Raptorial feeder | Herbivore | high | SRH |
| *P. trigla** | Small | Raptorial feeder | Herbivore | high | SRH |
| *P. vulgaris* | Small | Raptorial feeder | Herbivore | high | SRH |
| *Pompholyx* sp. | Small | Filter feeder | Detritivore | high | SFD |
| *P. complanate** | Small | Filter feeder | Detritivore | high | SFD |
| *P. sulcata* | Small | Filter feeder | Detritivore | high | SFD |
| *Proales* sp. | Small | Filter feeder | Detritivore | high | SFD |
| *Rotaria* sp. | Small | Filter feeder | Detritivore | high | SFD |
| *Schizocerca diversicornis* | Small | Filter feeder | Detritivore | high | SFD |
| *Synchaeta* sp.* | Small | Raptorial feeder | Herbivore | high | SRH |
| *S. stylata** | Small | Raptorial feeder | Herbivore | high | SRH |
| *S. tremula* | Small | Raptorial feeder | Herbivore | high | SRH |
| *Testudinella* sp. | Small | Filter feeder | Detritivore | high | SFD |
| *T. patina* | Small | Filter feeder | Detritivore | high | SFD |
| *Tetramastix opoliensis* | Small | Filter feeder | Detritivore | high | SFD |
| *Trichocerca* sp.* | Small | Raptorial feeder | Omnivore | high | SRO |
| *T. capucina** | Small | Raptorial feeder | Omnivore | high | SRO |
| *T. cylindrica** | Small | Raptorial feeder | Herbivore | high | SRH |
| *T. elongata* | Small | Raptorial feeder | Herbivore | high | SRH |
| *T. porcellus* | Small | Raptorial feeder | Omnivore | high | SRO |
| *T. similis* | Small | Raptorial feeder | Omnivore | high | SRO |
| *T. stylata* | Small | Raptorial feeder | Omnivore | high | SRO |
| *Trichotria tetractis* | Small | Filter feeding | Detritivore | high | SRO |

**Table S3.** Results of the principal coordination analysis showing the eigenvalue, viability, and cumulative variability of the first three coordinates, and the eigenvectors of the fish taxa for each principal coordination. The top 5 fish taxa with the highest absolute eigenvector value in these coordination axes are shown in bold.

| **Axis** | **PC1** | **PC2** | **PC3** |
|---|---|---|---|
| **Eigenvalue** | 33.04 | 15.86 | 11.32 |
| **Contribution (%)** | 17.58 | 8.44 | 6.02 |
| **Cumulative contribution (%)** | 17.58 | 26.02 | 32.04 |
| *Cobitis biwae* | 0.402 | 0.231 | 1.123 |
| *Cottus pollux* | -0.431 | -0.094 | 1.424 |
| *Cyprinus carpio* | 1.138 | -0.285 | **2.414** |
| *Garassius cuvieri* | 0.817 | 0.862 | 1.822 |
| *Garassius* sp. | 1.246 | -1.243 | 1.672 |
| *Gnathopogon* sp. | 0.802 | 0.942 | 1.431 |
| *Gymnogobius* sp. | 0.769 | 0.457 | -0.272 |
| *Hemibarbus labeo barbus* | 1.541 | **1.463** | 0.161 |
| *Hypomesus nipponiensis* | -0.203 | 0.753 | **3.140** |
| *Lepomis macrochirus* | 1.907 | 0.948 | -1.269 |
| *Leuciscus hakonensis / ezoensis* | -0.664 | **1.578** | 1.749 |
| *Micropterus salmoides salmoides* | **1.979** | 1.161 | -0.529 |
| *Misgurmus anguillicaudatus* | 0.212 | 0.405 | **2.576** |
| *Nipponocypris temminckii sieboldii* | 1.979 | 0.868 | -0.793 |
| *Oncorhynchus masou masou* | -0.780 | 0.186 | **2.829** |
| *Oncorhynchus mykiss* | -1.260 | 0.945 | 1.008 |
| *Opsariichthys uncirostris uncirostris* | 1.873 | **1.354** | -0.983 |
| *Phoxinus* sp. | 0.401 | 0.921 | **2.664** |
| *Plecoglossus altvelis altvelis* | 1.931 | 1.290 | -0.626 |
| *Pseudobagrus fulidraco* | 1.338 | 0.967 | -1.036 |
| *Pseudogobio ecocinus ecocinus* | **2.038** | 1.166 | -0.012 |
| *Pseudorasbora parva* | 0.724 | 0.558 | 2.328 |
| *Rhinogobius* sp. | **2.002** | -1.339 | 0.693 |
| *Salvelinus* sp. | **-2.296** | 1.095 | 0.889 |
| *Silurus asotus* | 1.762 | 1.219 | -0.311 |
| *Squalidus* sp. | 1.899 | **1.380** | -0.909 |
| *Tridentiger luroiwae breispinis* | 1.417 | 1.123 | -0.437 |
| *Zacco platypus* | **2.331** | **1.738** | 0.405 |

**Table S4.** Results of partial-dbRDA showing effects of environmental conditions and spatial configurations on taxa- and trait-based community structures.

| | taxa-based | | | trai-based | | |
|---|---|---|---|---|---|---|
| | **df** | **F** | **p value** | **df** | **F** | **p value** |
| Environment | 13 | 1.813 | 0.001 | 13 | 2.551 | 0.001 |
| Space | 13 | 1.768 | 0.001 | 13 | 2.049 | 0.001 |

**Table S5.** Results of variation partitioning of environmental variables (Env) and spatial variables (Spa) for taxa-based and trait-based community structures.

| Fraction | taxa-based | | trait-based | |
|---|---|---|---|---|
| | df | Adj R2 | df | Adj R2 |
| Total Env | 13 | 0.188 | 13 | 0.270 |
| Total Spa | 13 | 0.181 | 13 | 0.193 |
| Unique Env | 13 | 0.109 | 13 | 0.199 |
| Unique Spa | 13 | 0.102 | 13 | 0.122 |
| Env + Spa | 0 | 0.079 | 0 | 0.071 |
| Residual | 0 | 0.710 | 0 | 0.608 |

**Table S6.** Results of partial-dbRDA showing the association of species in the taxon-based community with significant environmental variables as determined by the permutation test. The top part of the table shows the coefficient of explanation (Adj-R2), the coefficient for axis 1, and the biplot score for each of the significant spatial variables, while the bottom part shows the eigenvector of each taxon for axis 1. For each variable, the top six taxa in terms of the absolute value of the eigenvector are indicated in bold.

| **Variables** | **WEF (PC1)** | **SDF (PC3)** | **Watershed** | **Volume** | **Height** |
|---|---|---|---|---|---|
| **Adj-R2** | 0.017 | 0.013 | 0.036 | 0.025 | 0.029 |
| **coefficient to axis1** | 0.407 | -0.473 | 0.124 | 0.115 | -0.138 |
| **biplot score** | 0.427 | -0.629 | 0.689 | 0.738 | -0.616 |
| *Alona* spp. | -0.158 | 0.157 | -0.052 | -0.118 | -0.288 |
| *Bosmina longilostris* | 0.097 | -0.055 | -0.286 | **-0.315** | -0.172 |
| *Bosminopsis deitersi* | -0.128 | -0.250 | 0.036 | -0.166 | 0.144 |
| *Ceriodaphnia* sp. | 0.024 | -0.013 | -0.131 | -0.039 | -0.097 |
| *Chydorus sphaericus* | -0.105 | 0.116 | 0.115 | **0.502** | 0.115 |
| *Daphnia galeata* | **0.329** | -0.058 | **-0.375** | -0.147 | **-0.374** |
| *Daphnia dentifera* | 0.156 | 0.143 | -0.159 | -0.066 | -0.087 |
| *Diaphanosoma brachyurum* | 0.127 | -0.009 | -0.184 | 0.046 | -0.119 |
| *Calanoida* | **0.308** | **0.217** | -0.291 | -0.024 | 0.017 |
| *Cyclopidae* | 0.032 | 0.193 | -0.094 | -0.193 | **-0.336** |
| *Cyclops vicinus* | **-0.415** | -0.150 | **-0.350** | 0.077 | -0.079 |
| *Eodiaptomus japonicus* | **0.238** | **0.231** | **-0.336** | 0.089 | -0.069 |
| *Ascomorpha ovalis* | 0.021 | **-0.333** | -0.221 | **-0.348** | -0.333 |
| *Ascomorpha* spp. | -0.011 | -0.006 | -0.250 | **0.117** | -0.031 |
| *Asplanchna priodonta* | -0.165 | -0.020 | -0.094 | 0.111 | **0.388** |
| *Asplanchna* sp. | **0.262** | -0.120 | -0.251 | -0.134 | -0.155 |
| *Cephalodella* sp. | -0.009 | 0.129 | 0.159 | -0.026 | 0.015 |
| *Collotheca* spp. | 0.021 | **-0.333** | -0.221 | **-0.348** | -0.333 |
| *Colurella* sp. | 0.027 | -0.117 | -0.031 | -0.266 | -0.158 |
| *Conochiloides* spp. | 0.202 | -0.107 | **-0.407** | -0.185 | -0.007 |
| *Conochilus* spp. | **0.351** | 0.041 | **-0.399** | -0.163 | **-0.381** |
| *Euchlanis dilatata* | -0.048 | 0.064 | **0.244** | -0.111 | 0.036 |
| *Filinia longiseta* | 0.217 | -0.034 | -0.208 | -0.233 | -0.009 |
| *Hexarthra mira* | -0.028 | 0.159 | -0.098 | 0.011 | -0.079 |
| *Kellicottia longispina* | **0.323** | 0.142 | -0.173 | 0.019 | 0.008 |
| *Keratella cochlearis* | 0.052 | -0.004 | -0.323 | -0.124 | -0.005 |
| *Keratella quadrata* | 0.031 | 0.124 | -0.334 | -0.131 | -0.293 |
| *Philodinidae* | -0.088 | 0.099 | 0.134 | -0.126 | -0.094 |
| *Ploesoma hudsoni* | 0.036 | **-0.286** | -0.141 | **-0.314** | -0.162 |
| *Ploesoma truncatum* | 0.188 | -0.041 | 0.109 | 0.162 | 0.147 |
| *Polyarthra euryptera* | 0.013 | 0.031 | -0.164 | -0.078 | 0.066 |
| *Polyarthra trigla* | -0.068 | 0.015 | -0.324 | -0.126 | -0.004 |
| *Pompholyx complanata* | -0.090 | -0.109 | -0.192 | 0.015 | 0.120 |
| *Synchaeta* sp. | 0.040 | -0.120 | -0.033 | -0.238 | **-0.441** |
| *Synchaeta stylata* | -0.049 | 0.040 | 0.012 | 0.031 | 0.318 |
| *Trichocerca capucina* | 0.130 | 0.076 | -0.220 | -0.001 | 0.098 |
| *Trichocerca cylindrica* | 0.157 | -0.007 | -0.213 | -0.095 | 0.031 |
| *Trichocerca* sp. | 0.028 | **-0.286** | -0.078 | **-0.428** | **-0.352** |

**Table S7.** Results of partial-dbRDA showing the association of species in the taxon-based community with significant spatial variables as determined by the permutation test. The top part of the table shows the coefficient of explanation (Adj-R2), the coefficient for axis 1, and the biplot score for each of the significant spatial variables, while the bottom part shows the eigenvector of each taxon for axis 1. For each variable, the top six taxa in terms of the absolute value of the eigenvector are indicated in bold.

| **Variables** | **Elevation** | **MEM2** | **MEM3** | **MEM4** | **MEM5** | **MEM9** | **MEM10** | **MEM11** |
|---|---|---|---|---|---|---|---|---|
| **Adj-$R^2$** | 0.020 | 0.020 | 0.021 | 0.015 | 0.016 | 0.022 | 0.014 | 0.015 |
| **coefficient to axis1** | 0.144 | 0.143 | 0.191 | 0.102 | 0.100 | 0.091 | -0.100 | 0.092 |
| **biplot score** | 0.589 | 0.593 | 0.445 | 0.832 | 0.850 | 0.932 | -0.851 | 0.924 |
| *Alona* spp. | -0.215 | -0.121 | 0.124 | -0.062 | 0.198 | -0.004 | -0.105 | **-0.291** |
| *Bosmina longilostris* | 0.008 | -0.032 | 0.200 | -0.103 | -0.019 | 0.122 | 0.079 | -0.183 |
| *Bosminopsis deitersi* | -0.036 | 0.175 | -0.166 | -0.094 | -0.122 | **-0.322** | **0.224** | **-0.348** |
| *Ceriodaphnia* sp. | 0.016 | 0.118 | 0.059 | 0.078 | **-0.251** | 0.218 | 0.075 | 0.022 |
| *Chydorus sphaericus* | -0.004 | -0.034 | 0.083 | **0.395** | 0.121 | 0.069 | -0.099 | **-0.266** |
| *Daphnia galeata* | 0.056 | 0.140 | 0.054 | -0.011 | 0.006 | 0.128 | **-0.374** | **-0.276** |
| *Daphnia dentifera* | 0.178 | -0.085 | 0.122 | 0.006 | 0.175 | 0.234 | **-0.455** | -0.195 |
| *Diaphanosoma brachyurum* | -0.096 | 0.121 | -0.239 | 0.070 | -0.048 | 0.090 | -0.011 | 0.027 |
| *Calanoida* | -0.062 | -0.153 | 0.088 | 0.018 | 0.027 | **0.304** | 0.011 | 0.054 |
| *Cyclopidae* | **-0.256** | -0.081 | -0.048 | -0.113 | 0.008 | -0.033 | 0.123 | -0.027 |
| *Cyclops vicinus* | **-0.635** | -0.086 | 0.227 | **-0.339** | -0.051 | -0.156 | -0.068 | 0.113 |
| *Eodiaptomus japonicus* | -0.067 | **-0.221** | 0.072 | 0.004 | **-0.253** | 0.118 | -0.174 | 0.046 |
| *Ascomorpha ovalis* | -0.055 | **0.381** | -0.096 | 0.063 | -0.175 | **0.296** | -0.015 | 0.160 |
| *Ascomorpha* spp. | **-0.280** | -0.078 | -0.046 | -0.139 | -0.016 | 0.035 | 0.128 | 0.127 |
| *Asplanchna priodonta* | 0.064 | 0.041 | **0.320** | -0.072 | -0.111 | -0.295 | 0.062 | -0.230 |
| *Asplanchna* sp. | -0.193 | -0.079 | **-0.254** | -0.060 | 0.077 | **0.367** | 0.126 | **0.278** |
| *Cephalodella* sp. | -0.254 | **-0.328** | 0.235 | -0.147 | **0.254** | 0.026 | 0.079 | -0.001 |
| *Collotheca* spp. | -0.055 | **0.381** | -0.096 | 0.063 | -0.175 | **0.296** | -0.015 | 0.160 |
| *Colurella* sp. | -0.024 | **0.317** | -0.232 | -0.121 | -0.198 | -0.040 | 0.002 | 0.062 |
| *Conochiloides* spp. | -0.117 | 0.189 | **-0.259** | 0.108 | -0.173 | -0.030 | 0.007 | 0.111 |
| *Conochilus* spp. | **-0.262** | -0.103 | 0.129 | 0.048 | -0.047 | 0.257 | 0.080 | -0.012 |
| *Euchlanis dilatata* | -0.159 | 0.000 | -0.066 | 0.059 | 0.112 | -0.039 | 0.167 | -0.203 |
| *Filinia longiseta* | 0.114 | -0.036 | 0.235 | -0.057 | 0.197 | **0.346** | -0.117 | -0.183 |
| *Hexarthra mira* | 0.014 | -0.075 | **0.266** | **0.306** | -0.202 | -0.214 | 0.115 | -0.002 |
| *Kellicottia longispina* | -0.035 | -0.087 | 0.096 | 0.276 | **0.222** | 0.215 | **0.200** | -0.099 |
| *Keratella cochlearis* | -0.172 | -0.142 | 0.068 | 0.023 | 0.130 | -0.061 | -0.080 | -0.038 |
| *Keratella quadrata* | -0.111 | -0.208 | **0.245** | -0.012 | 0.035 | -0.060 | -0.042 | -0.047 |
| *Philodinidae* | -0.132 | -0.020 | -0.052 | -0.112 | 0.220 | 0.218 | 0.043 | **-0.311** |
| *Ploesoma hudsoni* | 0.069 | **0.301** | 0.232 | -0.005 | **-0.290** | 0.024 | 0.022 | 0.049 |
| *Ploesoma truncatum* | **0.367** | -0.060 | 0.216 | **0.291** | -0.089 | -0.219 | **0.325** | 0.147 |
| *Polyarthra euryptera* | -0.024 | 0.198 | -0.150 | -0.114 | -0.164 | 0.006 | -0.007 | 0.037 |
| *Polyarthra trigla* | -0.051 | 0.070 | -0.140 | 0.040 | **-0.329** | 0.131 | -0.148 | -0.031 |
| *Pompholyx complanata* | -0.037 | 0.175 | -0.167 | -0.047 | -0.206 | 0.009 | 0.171 | -0.033 |
| *Synchaeta* sp. | -0.110 | 0.102 | -0.183 | **-0.283** | 0.085 | 0.155 | 0.106 | 0.155 |
| *Synchaeta stylata* | 0.015 | 0.120 | 0.217 | **0.387** | 0.041 | -0.086 | -0.041 | -0.148 |
| *Trichocerca capucina* | 0.026 | 0.010 | 0.015 | 0.078 | -0.154 | 0.081 | -0.024 | 0.071 |
| *Trichocerca cylindrica* | -0.074 | 0.192 | **-0.297** | 0.040 | -0.121 | 0.170 | 0.126 | 0.065 |
| *Trichocerca* sp. | **-0.255** | **0.339** | **-0.241** | -0.083 | -0.117 | -0.098 | **0.252** | -0.033 |

**Table S8.** Results of partial-dbRDA showing the association of functional groups in the trait-based community with significant environmental variables as determined by the permutation test. The top part of the table shows the coefficient of explanation (Adj-R2), the coefficient for axis 1, and the biplot score for each of the significant spatial variables, while the bottom part shows the eigenvector of each functional group for axis 1. For each variable, the top six functional groups in terms of the absolute value of the eigenvector are shown in bold.

| Variables | WEF (PC1) | SDF (PC2) | Watershed | Volume | Height | Age |
|---|---|---|---|---|---|---|
| **Adj-R2** | 0.069 | 0.037 | 0.061 | 0.059 | 0.021 | 0.018 |
| **coefficient to axis1** | 0.407 | -0.654 | 0.124 | 0.115 | -0.138 | 0.118 |
| **biplot score** | 0.427 | -0.384 | 0.689 | 0.738 | -0.616 | 0.723 |
| LRC | -0.009 | 0.086 | **0.009** | **0.634** | **0.289** | **-0.053** |
| LFH | **0.633** | **-0.081** | **-0.374** | 0.151 | **-0.357** | **-0.330** |
| LBO | 0.185 | -0.016 | **-0.442** | 0.084 | **0.174** | 0.149 |
| LFO | **0.670** | **-0.213** | **-0.769** | **-0.110** | **-0.336** | **0.401** |
| MFH | **0.388** | -0.045 | -0.323 | **0.328** | 0.046 | 0.036 |
| MRO | **-0.029** | -0.032 | -0.084 | -0.025 | -0.155 | 0.042 |
| MRC | 0.185 | -0.016 | **-0.442** | 0.084 | **0.174** | 0.149 |
| SFH | **-0.290** | **0.249** | -0.054 | **0.318** | 0.010 | **0.158** |
| SFDH | -0.012 | -0.029 | -0.166 | **-0.510** | -0.109 | -0.008 |
| SRH | 0.068 | **-0.093** | **0.042** | **-0.427** | **-0.205** | **-0.021** |
| SRO | 0.028 | **0.652** | -0.226 | 0.221 | **0.085** | 0.144 |
| SFD | 0.073 | 0.140 | -0.033 | 0.004 | -0.139 | 0.150 |

**Table S9.** Results of partial-dbRDA showing the association of functional groups in the trait-based community with significant spatial variables determined by the permutation test. The top part of the table shows the coefficient of explanation (Adj-R2), the coefficient for axis 1, and the biplot score for each of the significant spatial variables, while the bottom part shows the eigenvector of each functional group for axis 1. For each variable, the top six functional groups in terms of the absolute value of the eigenvector are indicated in bold.

| **Variables** | **Elevation** | **MEM1** | **MEM3** | **MEM4** | **MEM9** | **MEM10** |
|---|---|---|---|---|---|---|
| **Adj-R2** | 0.030 | 0.039 | 0.020 | 0.028 | 0.043 | 0.051 |
| **coefficient to axis1** | 0.145 | -0.189 | 0.191 | 0.102 | 0.091 | -0.100 |
| **biplot score** | 0589 | -0.449 | 0.445 | 0.832 | 0.932 | -0.851 |
| LRC | **0.178** | 0.041 | 0.070 | -0.053 | 0.085 | **-0.319** |
| LFH | **0.525** | **-0.381** | **0.299** | **0.169** | **0.523** | **-0.833** |
| LBO | -0.232 | -0.028 | 0.078 | **-0.299** | **0.169** | **0.251** |
| LFO | -0.204 | **-0.570** | 0.098 | **-0.178** | **0.460** | **-0.290** |
| MFH | -0.244 | 0.177 | **-0.166** | -0.049 | 0.022 | **0.086** |
| MRO | -0.191 | -0.009 | -0.014 | **-0.264** | 0.077 | -0.079 |
| MRC | -0.232 | -0.028 | 0.078 | **-0.299** | **0.169** | **0.251** |
| SFH | -0.237 | **0.211** | 0.061 | 0.091 | **-0.524** | -0.155 |
| SFDH | -0.051 | -0.032 | **-0.018** | -0.055 | **-0.067** | **0.179** |
| SRH | 0.058 | **-0.135** | **-0.037** | -0.095 | 0.165 | -0.018 |
| SRO | **0.170** | **0.265** | **0.471** | **0.576** | **-0.211** | 0.080 |
| SFD | -0.084 | 0.049 | 0.089 | **0.101** | 0.036 | 0.079 |
